# Multilevel gene expression changes in lineages containing adaptive copy number variants

**DOI:** 10.1101/2023.10.20.563336

**Authors:** Pieter Spealman, Carolina de Santana, Titir De, David Gresham

## Abstract

Copy-number variants (CNVs) are an important class of recurrent variants that mediate adaptive evolution. While CNVs can increase the relative fitness of the organism, they can also incur a cost. We previously evolved populations of *Saccharomyces cerevisiae* over hundreds of generations in glutamine-limited (Gln-) chemostats and observed the recurrent evolution of CNVs at the *GAP1* locus. To understand the role that expression plays in adaptation, both in relation to the adaptation of the organism to the selective condition, and as a consequence of the CNV, we measured the transcriptome, translatome, and proteome of 4 strains of evolved yeast, each with a unique CNV, and their ancestor in Gln-conditions. We find CNV-amplified genes correlate with higher RNA abundance; however, this effect is reduced at the level of the proteome, consistent with post-transcriptional dosage compensation. By normalizing each level of expression by the abundance of the preceding step we were able to identify widespread divergence in the efficiency of each step in the gene in the efficiency of each step in gene expression. Genes with significantly different translational efficiency were enriched for potential regulatory mechanisms including either upstream open reading frames, RNA binding sites for SSD1, or both. Genes with lower protein expression efficiency were enriched for genes encoding proteins in protein complexes. Taken together, our study reveals widespread changes in gene expression at multiple regulatory levels in lineages containing adaptive CNVs highlighting the diverse ways in which adaptive evolution shapes gene expression.

## Introduction

Copy-number variants (CNVs) are amplifications or deletions of DNA that can span dozens of nucleotides to whole chromosomes. CNVs are frequently observed over both short and long evolutionary time spans, although their selective advantage may be different between the two (Kuzmin et al. 2022). In the short term CNVs can result in large changes in gene expression and protein abundance, which provides a selective advantage further driving rapid adaptive evolution (Kondrashov 2012; Myhre et al. 2013; Robinson et al. 2023). Gene amplification has been shown to mediate rapid adaptation to a variety of selective pressures from nutrient limitation to antibiotics in both natural and experimental populations of microbes (Hull et al. 2017; Gresham et al. 2008; Nair et al. 2008; Selmecki et al. 2009; Paulander et al. 2010; Pränting and Andersson 2011; Hong and Gresham 2014; Dhami et al. 2016; Lauer et al. 2018; Todd and Selmecki 2020). CNVs are also common in cancers, where they can promote tumorigenesis by oncogene amplification (Ben-David and Amon 2020) and drive genome-wide changes in gene expression (Shao et al. 2019).

A simple model of the short-term fitness effects of CNVs is a variant of the “driver-hitchhiker” model (Smith and Haigh 1974; Golzio and Katsanis 2013) wherein the fitness benefit of a CNV is derived from the amplification of a single gene or “driver”, whereas fitness costs arise from the amplification of “hitchhikers”. Since CNVs include up to hundreds of genes, the potential fitness costs of hitchhikers can be high (Makanae et al. 2013; Avecilla et al. 2023). These fitness costs can be categorized as “dosage burden” wherein the fitness cost arises from the burden of the additional DNA replication and gene expression (Makanae et al. 2013; Dekel and Alon 2005; Bonney et al. 2015). Conversely, fitness costs may arise from the stoichiometric imbalance of specific “dosage sensitive” genes. The archetypal example of dosage sensitivity is the imbalance of proteins involved in a heteromeric protein complex, leading to negative impacts on complex formation and function (Rice and McLysaght 2017; Wagner 2005; Veitia et al. 2008). Importantly, the fitness effect of any CNV is highly dependent on the genetic background and environmental context (Robinson et al. 2023; Sunshine et al. 2015).

Dosage compensation (DC) is one mechanism by which an organism could mitigate the fitness costs of hitchhiker gene amplification. While classically associated with sex chromosome inactivation through chromatin silencing (Straub and Becker 2007; Jordan et al. 2019), DC can also be gene specific (Straub and Becker 2007). Furthermore, while DC is often conceptualized as ‘complete’, such that additional gene copies result in no additional expression, DC can also result in a range of increased gene expression levels that are significantly less than what would be expected based on the copy-number (Itoh et al. 2007; Brockdorff and Turner 2015). This ‘incomplete’ dosage compensation is sometimes referred to as attenuation (Ascencio et al. 2021). Here, we use the term dosage compensation (DC) to refer to both ‘complete’ (e.g, the gene expression from two copies of the gene is identical to the gene expression from one copy) and ‘incomplete’ (e.g., the gene expression from two copies of the gene is greater than the gene expression from one copy, but less than two times the expression of one copy).

Recent work on the transcriptional regulation of CNV amplified genes has found that, depending on genetic background and environment 10-60% of amplified genes show evidence of transcriptional dosage compensation (Avecilla et al. 2023; Springer et al. 2010; Gasch et al. 2016). The mechanism underlying this DC is unclear but changes in transcription factor abundances and their concentrations is one possible mechanism (Prestel et al. 2010).

Translational DC has recently become a focus of interest with the advent of ribosome profiling (Ingolia et al. 2009; Zhang and Presgraves 2017) but has also been studied using other methods (Gebauer et al. 1999; Krasovec et al. 2023). At the translational level some global mechanisms have been proposed including spliceosomal or ribosomal components acting to limit total global expression in an organism (Veitia et al. 2013), and co-translational mechanisms, such as the unfolded protein response (Kane et al. 2021), although the universality of these mechanisms remains unresolved (Larrimore et al. 2020). Gene specific mechanisms have also been proposed and identified, such as sequence specific RNA-binding proteins (RBPs) acting on specific genes and altering their post-transcriptional and translational regulation (Mayr 2017; Schultz et al. 2020). The archetypal case of translational DC is sex-chromosome dosage compensation in *Drosophila melanogaster*, enacted by the RNA-binding protein *SXL* altering the efficacy of alternative translation initiation at an upstream open reading frame (uORF) (Medenbach et al. 2011).

uORFs are short ORFs (6-300 nucleotides long) that are positioned upstream, in the transcript leader (TL, or 5’ UTR) of the main ORF (mORF) of a gene. uORFs have either canonical start codons (e.g. AUG) or near-cognate codon (NCC, (Zitomer et al. 1984)) translation initiation sites (TIS) that allow for alternative translation initiation separate from the mORF, and as such can operate as translational regulatory elements, either preventing or enabling the translation of the downstream mORF (Hinnebusch 2005). While uORFs have been suggested to be a broad class of stress responsive regulatory elements (Lawless et al. 2009) their role in the CNVs has not previously been evaluated.

One interesting candidate for gene specific translational regulation in *S. cerevisiae* is *SSD1*, which encodes an RBP found to preferentially bind to cis-elements in transcript leaders (TLs) (Bayne et al. 2022) and alter translation (Hogan et al. 2008; Ohyama et al. 2010). SSD1 is believed to play an important role in the response of yeast to aneuploid stress (Hose et al. 2015; Hose et al. 2020). While initial research suggested that yeast exhibit high sensitivity to aneuploidy, subsequent research has shown that this sensitivity is strain dependent, and due to the loss of *SSD1* in the lab strain W303 (Hose et al. 2015; Hose et al. 2020). Whether *SSD1* has a role in accommodating CNVs in the genome is unknown.

Quantitative mass spectrometry has enabled the identification of DC at the level of protein abundance (Stingele et al. 2012; Senger et al. 2022). At the protein level, work in yeast has suggested 10-20% of gene amplifications have significant DC (Dephoure et al. 2014; Ishikawa et al. 2017), although genetic differences are a known source of variance (Larrimore et al. 2020; Hose et al. 2020). One proposed mechanism for DC at the protein level is that excess subunits of proteins associated with heteromeric complexes are selectively targeted for protein aggregation (Samant et al. 2019) and degradation by ubiquitination (Senger et al. 2022; Ishikawa et al. 2020). A recent study using hundreds of natural isolates reported that nearly all aneuploid yeast strains studied had consistent protein level DC with an average decrease of 25% from the expected abundance (Muenzner et al. 2022).

Here, we undertook an analysis of multi-level gene expression changes in long-term experimentally evolved lineages of *Saccharomyces cerevisiae* with acquired adaptive CNVs. These strains evolved over the course of hundreds of generations in glutamine-limited chemostats and contain distinct adaptive CNVs containing the same driver gene *GAP1* (Lauer et al. 2018) but distinct sets of additional hitchhiker genes within the CNV, numbering between 18 to 82. We quantified expression at the transcriptional, translational, and protein levels in the glutamine-limited chemostats. We found over 2,000 genes were significantly different in transcript abundances and ribosome occupancy, and over 1700 genes were significantly different in protein abundance.

By normalizing each step of gene expression to account for changes at the preceding level, we were able to identify changes in gene regulation that alter the efficiency, including both increases and decreases, of expression at each level. We find evidence of widespread changes in gene expression efficiency at the transcriptional (median 25%), translational (median 11%), and protein (median 4%) levels. Decreased expression efficiency is one possible mechanism of dosage compensation of CNV-amplified genes and we find evidence of significantly decreased expression efficiency at the transcriptional (median 21%), translational (median 7%), and protein abundance (median 9%) levels specifically for CNV-amplified genes. We find CNV amplified genes have higher rates of genes with lower expression efficiency at every level of expression. However, it is only at the level of protein expression efficiency this is significantly different from the global rate.

We find that many genes with significant differences in expression efficiency are also enriched for potential regulatory mechanisms. Genes with significantly different translation efficiencies are enriched in both uORFs and *SSD1* binding site motifs, as well as the co-occurrence of uORFs and *SSD1* binding sites. Genes with significantly different protein expression efficiency are also significantly enriched in genes associated with protein complexes, suggesting stoichiometric imbalances may lead to targeted protein degradation.

Notably, each mechanism that we identify as contributing to dosage compensation is also involved in the regulation of gene expression more broadly and is not specific to CNV amplified genes. This indicates that dosage compensation mechanisms in yeast that act on additional copies of genes are not unique mechanisms but rather, extensions of the suite of gene regulatory mechanisms that act to maintain a robust phenotype in response to diverse cellular and environmental stresses.

## Results

### Topological structure of copy number variant alleles

We isolated four lineages that contained CNVs at the *GAP1* locus following long term selection in glutamine-limited chemostats. CNV-containing lineages were identified through the use of a CNV reporter system (Lauer et al. 2018; Spealman et al. 2023) and isolated using FACS from heterogeneous populations following more than 150 generations of selection (**Fig 1A**). Defining the gene expression consequences of altered copy number requires precise definition of the copy number of each gene. Therefore, we performed hybrid *de novo* genome assembly, using a combination of long-read (Oxford Nanopore Technologies) and short-read (Illumina) sequencing to determine the topological structures for the CNVs in each strain (**Fig 1B**). On the basis of the CNV structures we also inferred the likely mechanisms of formation, which include transposon mediated tandem amplification and origin of replication dependent amplification (ODIRA, (Brewer et al. 2011)) and named the CNV alleles to reflect these mechanisms. Because of the difficulty in ONT sequencing of inverted repeat sequences (Spealman et al. 2020) all ODIRA events were fully resolved using manual analysis.

**Fig 1.**
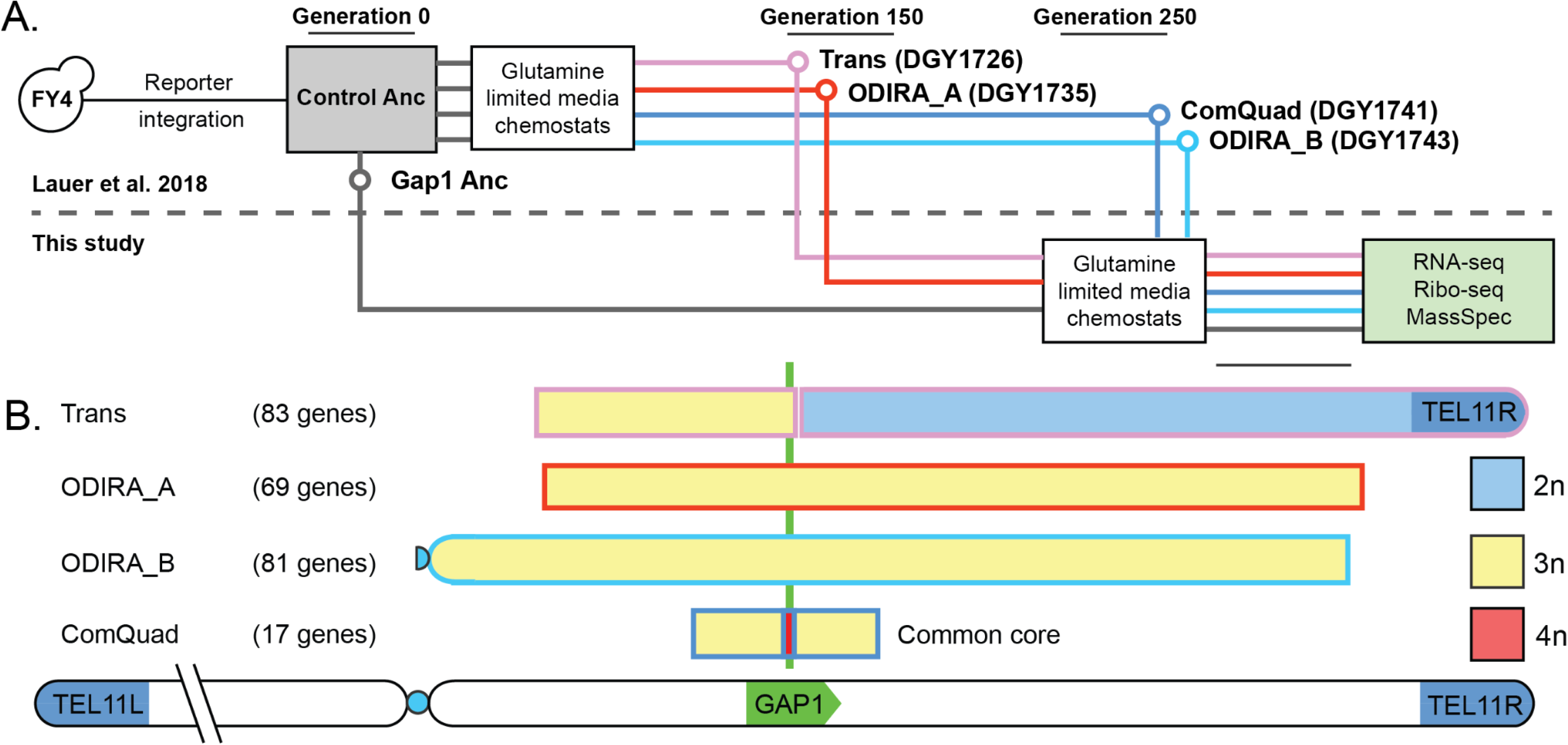
Study design and characterization of CNV strains. (**A**) Strain provenance ((Lauer et al. 2018), above the dotted line) and experimental design to assess gene expression effects using RNA-seq, ribosome profiling (Ribo-seq), and TMT-labeled mass spectrometry (this study, beneath dotted line) in glutamine-limited chemostats. (**B**) A schematic showing the *GAP1*-locus amplified genes in each strain and their copy-number. A common core of 17 genes is amplified in every evolved strain.

CNV structures include “Trans”, a transposon mediated tandem duplication, spanning the genes from *TOF2* to *GAP1*, nested within a translocation and telomere replacement on ChrVIII (Figure SF1C). “ODIRA_A”, an ODIRA event triplication spanning 77 genes (*ALY1*-*SIR1*) (Figure SF1D) and “ODIRA_B”, an ODIRA event triplication spanning 91 genes (*VPS1*-*ESL2*) (Figure SF1F). Finally, a complex quadruplex or “ComQuad’’ resulting from a ODIRA driven triplication followed by a singular amplification of *GAP1* by a transposon event that spans the common core 17 genes from (*SET3*-*TRK2*) (Figure SF1D). Using the method we unambiguously determined the copy number of each gene in each CNV (**Table ST1**). Collectively, the four CNV strains include between 17 – 91 genes present at 2, 3, or 4 additional copies. A set of 17 amplified genes centered on the *GAP1* gene is common to all strains. A median of two *de novo* SNVs or indels were also identified per strain (**Figure SF2**).

To assess the gene expression consequences of both CNVs and adaptation to nutrient limitation, we analyzed the four evolved CNV strains and the ancestral wildtype in glutamine-limited chemostats. Strains were grown until reaching a steady-state condition during which culture density remained constant (∼10-12 generations). Cycloheximide (100μg/L) was added to cells for 2 minutes before harvesting by vacuum filtration. All RNA-seq and ribosome profiling experiments were performed in tandem using two biological replicates for each strain. Mass spectrometry was conducted using 5 biological replicates to avoid high false detection rates (Martinez-Val et al. 2016; Brenes et al. 2019).

### Divergence of gene expression at multiple levels in evolved strains

We first sought to compare gene expression between each evolved strain and the ancestor. We retained 4,291 after filtering genes poorly represented in any strain at any level of expression. We found 2,371 genes were significantly different in RNA abundance in at least one evolved strain (DESeq2, Benjamini-Hochberg (BH) adjusted p-value < 0.05, **Table ST4**). Similarly, 2,079 genes were significantly different in RPF abundance (DESeq2, BH adj.p-value < 0.05, **Table ST5**). Finally, 1,785 genes were significantly different in MS abundance (Welch’s t-test, BH adj.p-value < 0.05, **Table ST6**). As the scale and nature of the three different datatypes differ, we performed a unit transformation to enable direct comparison between assays (**methods**). To visualize gene expression across multiple levels we made a heatmap using the divergence in expression between each evolved and the ancestor strain. (**Fig. 2A, Figure SF3**). We used k-means clustering on all levels of expression across all evolved strains (**Table ST7**) and found some clusters are significantly enriched in genes associated with specific gene ontology terms (SGD, BH adj.p-value < 0.05). For example clusters 3, 4, and 5 which trend lower in expression in the evolved strains are significantly enriched in genes associated with gene ontologies. Cluster 3 is enriched in ‘protein catabolic process’ (GO:0030163); cluster 4 with ‘protein-containing complex organization’ (GO:0043933); and cluster 5 with ‘energy derivation by oxidation of organic compounds’ (GO:0015980). Notably, clusters 2, 13, and 15 which tend to be higher in expression in the evolved strains, were not enriched in any GO terms; however they were significantly enriched in genes amplified by the CNV (FET, p-value < 0.05).

**Fig 2.**
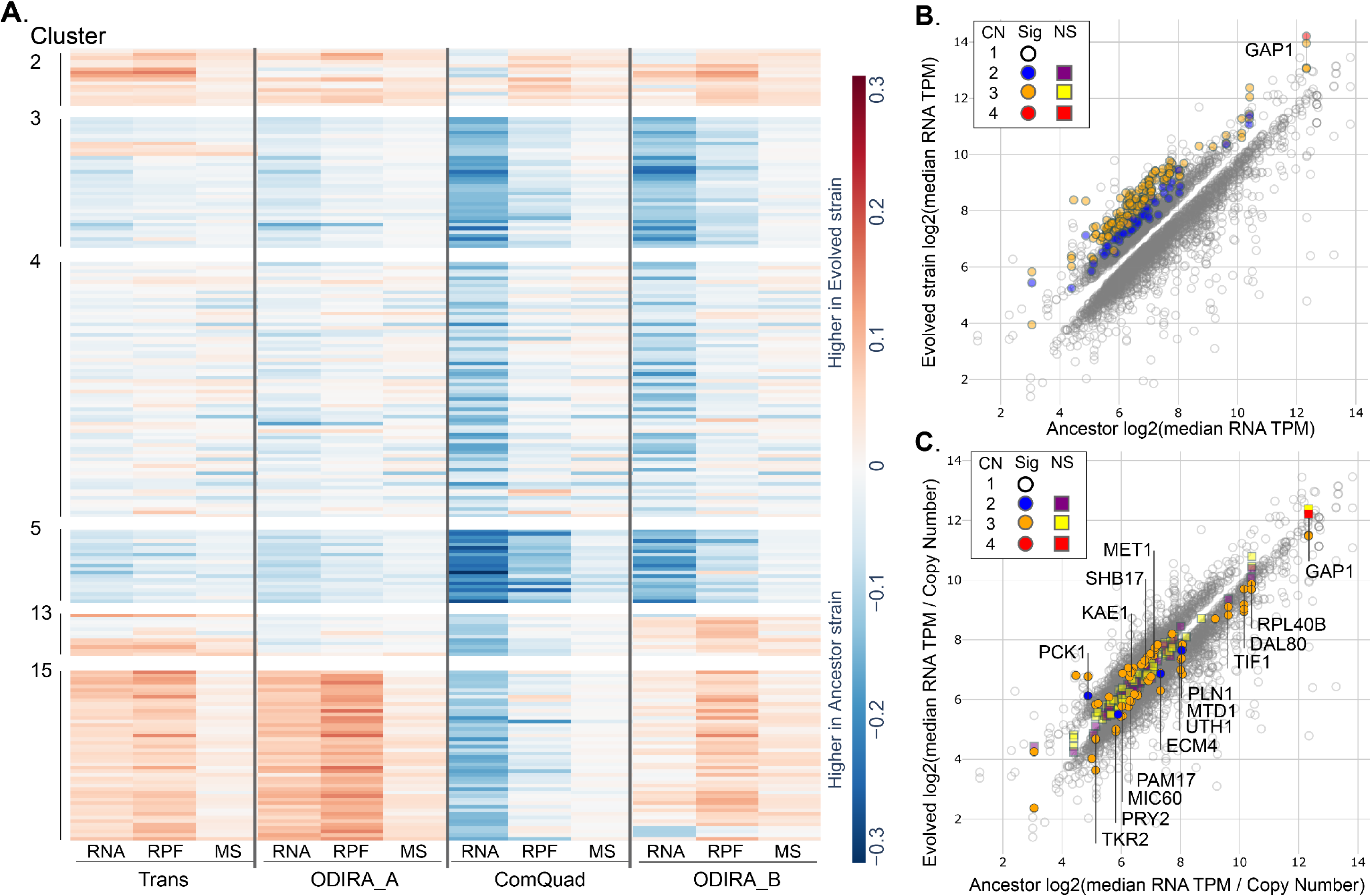
Gene expression divergence in evolved lineages containing CNVs. (**A**) A heatmap of relative unit normalized (See methods) expression at multiple levels (RNA, RPF, MS) of genes from all strains (complete heatmap available as **Figure SF3**). The 218 genes shown are all significantly different in abundance (DESeq2 (RNA, RPF); Welch’s t-test (MS); BH adj. P-value < 0.01) at multiple levels of gene expression, or are CNV amplified in at least on strain. (**B**) Scatterplot of RNA abundance of genes with significantly different (DESeq2, BH adj. P-value < 0.05) expression between each of the four evolved strains and the ancestor. All CNV amplified genes are significantly higher in the evolved strain. (**C**) To assess if there were changes in transcription efficiency we corrected RNA abundance for gene copy number (CN) and found that a minority (median of 32%) of genes exhibited significantly different expression than expected (DESeq2, BH adj.p-value < 0.05).

### Differences in transcription efficiency in CNV amplified genes

Whereas all CNV amplified genes were found to have higher RNA abundance than the ancestor (**Fig. 2B**), previous work has described gene specific transcription dosage compensation of CNV amplified genes (Straub and Becker 2007; Ascencio et al. 2021). Typically, in differential expression analysis the copy-number of the underlying genes are assumed to be uniform throughout the genome and between genotypes. As that is not the case in our study, we first sought to correct for changes in copy-number. Here, we define a null model in which the expected RNA abundance for genes within a CNV is determined by the abundance in the ancestor multiplied by the copy-number in the evolved strain. For example, a two fold increase in gene copy-number is expected to result in a two-fold increase in mRNA. We determined the copy number corrected gene expression for all genes and then compared the observed RNA abundance to this expected value to quantify differences in transcript efficiency (**Table ST8**), (**Fig 2C**).

In total, almost half (46 of 91) of the CNV amplified genes were found to have significantly different RNA abundances from what would be expected given their copy-number. We found significantly increased transcription efficiency for 21 (23%) CNV amplified genes, with 4 genes (*PCK1*, *KAE1*, *SHB17*, and *MET1*) being significantly higher in multiple lineages, suggesting increased transcription efficiency. Conversely, we found significantly lower RNA transcription efficiency in 24 (26%) genes, with 12 genes showing decreased transcription efficiency in all strains (*TKR2*, *PRY2*, *MIC60*, *PAM17*, *ECM4*, *UTH1*, *MTD1*, *PLN1*, *TIF1*, *DAL80*, *RPL40B*, and *GAP1*). Interestingly, we find that *GAP1*, the hypothesized driver of CNV selection, has lower transcription efficiency in all evolved strains, although this reduction is only significant in two of the CNV strains (Trans and ODIRA_A). *DAL80*, a transcription factor that is the negative regulator of *GAP1* (Daugherty et al. 1993), has significantly lower transcription efficiency in all evolved strains.

While this supports the model of transcription efficiency as a mechanism of dosage compensation, decreased transcription efficiency is also prevalent in non-amplified genes. Indeed, the frequency of non-amplified genes with significant divergence in transcript efficiency is very similar to the frequency observed in CNV amplified genes with a median 898 genes per strain (21%). This suggests that the decreased transcription efficiency observed in CNVs may not be produced by a mechanism unique to them.

### Changes in translation efficiency

To investigate how changes in translational regulation may act to alter gene expression, we normalized the ribosome abundance, as measured by RPFs aligned to mORFs only, by transcript abundance (**Table ST9**). This translation efficiency ratio is an estimate of the amount of translating ribosomes per transcript (Ingolia et al. 2009) and can be the result of altered translation initiation rates at the start codon as well as differences in elongation rates throughout the mORF (Weinberg et al. 2016).

In total, 22 CNV amplified genes were found to have significantly different translation efficiencies (FET, p-value < 0.05). Of these, 9 had significantly higher translation efficiency in the evolved strain relative to the ancestor with 5 genes (*PLN1*, *PAM17*, *MTD1*, *RPL40B*, and *DID2*) being higher in multiple backgrounds. Conversely, 10 genes had significantly lower translation efficiency relative to the ancestor but only two were significantly lower in multiple backgrounds: *TIF1* in both Trans and ODIRA_A, as well as *GAP1* in every background (**Table ST9**). Finally, 3 genes were differentially expressed in specific strains, *DAL80* was higher in Trans and lower in ODIRA_B; *RPS21A* was higher is ODIRA_A but lower in both Trans and ODIRA_B; while *UTH1* was lower in ODIRA_A and higher in both ODIRA_B and ComQuad.

For the unamplified genes, we found that 1011 genes had significantly different translation efficiencies relative to the ancestor (**Fig 3A**, FET, p-value < 0.05). The majority (655, 65%) had significantly higher translation efficiency than the ancestor, with 371 genes being higher in multiple backgrounds, suggesting they are translationally up-regulated. Conversely, only 279 (28%) of these had significantly lower translation efficiency than the ancestor, with 100 being lower in multiple backgrounds. Finally, 77 genes (8%) were significantly different in opposite directions in specific strains.

**Fig 3.**
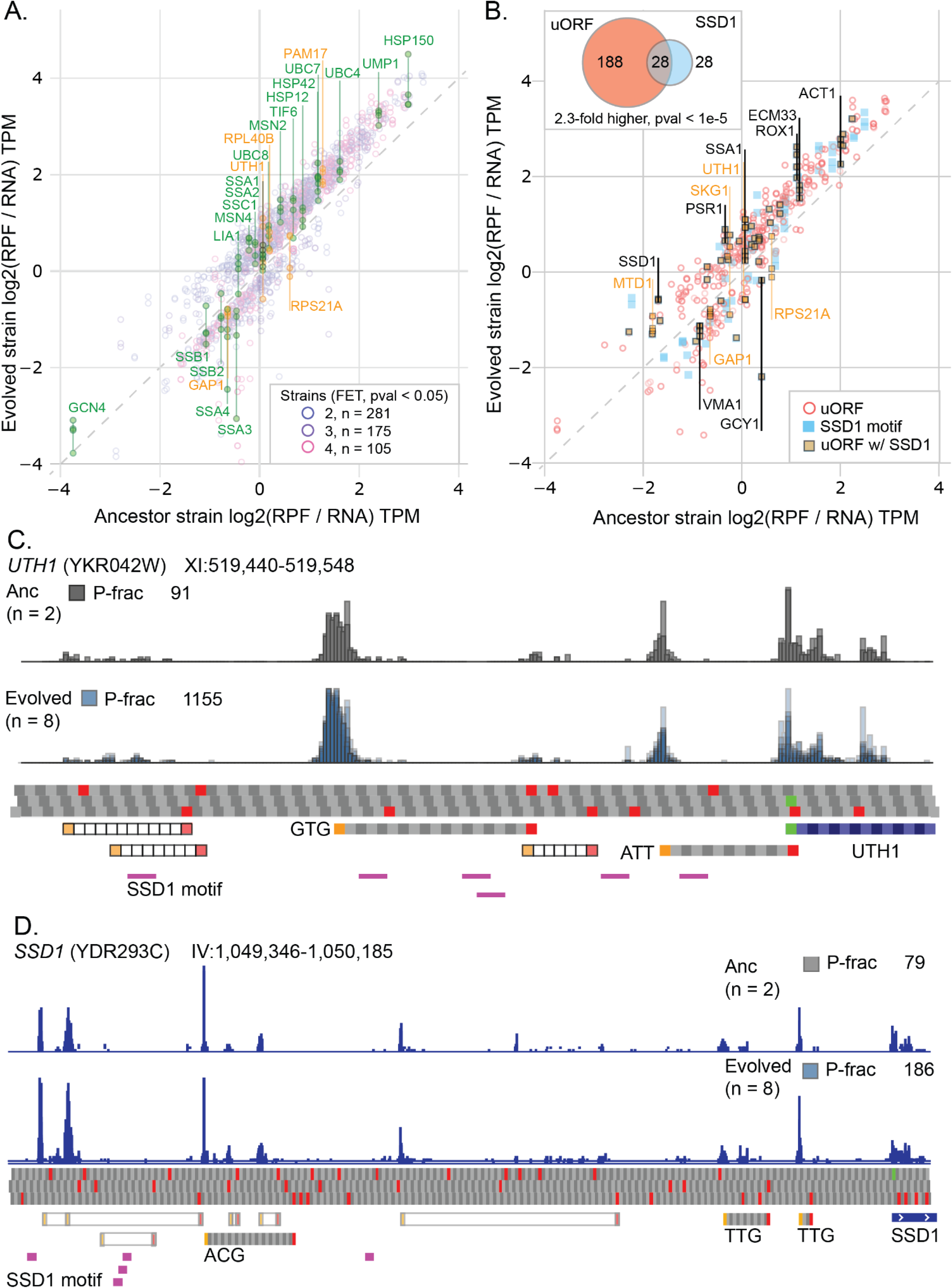
– Divergence in translation efficiencies and potential regulatory mechanisms. (**A**) 561 genes with significantly different translation efficiency relative to the ancestor in at least two evolved strains (FET, p-value < 0.05). Gene names in orange indicate CNV amplified genes. (**B**) A subset of genes with significantly different translation efficiency and the presence of a high confidence uORF (red), significant SSD1 motifs (blue), or both (2.3-fold higher than expected at random, HGM, p-value < 1e-5). (**C**) Ribosome pile-ups upstream of *UTH1* in the ancestor and evolved strains. High confidence predicted uORFs are shown in gray codons, low confidence in hollow boxes. *SSD1* binding motifs are shown in purple. (**D**). Example of uORFs and *SSD1* binding motifs upstream of *SSD1*.

We find that genes amplified in CNVs more frequently have lower translation efficiencies than the genome background in three of the four strains, although only weakly significant. For example, in Trans, 5 of the 85 (6.0%) genes amplified in CNVs have significantly lower translation efficiency. This is 2.3-fold higher that the rate observed in the rest of the genome where 112 of the 4,206 (2.7%) unamplified genes have significantly lower translation efficiency observed (FET, p-value = 0.08). Similarly, ODIRA_A with 4 of 69 (5.8%) is 2.9-fold higher than the background (2.2%), (FET, p-value = 0.06), ComQuad with 2 of 17 (11.2%) is 4.7-fold higher than the background (3.1%), (FET, p-value = 0.08). While ODIRA_B, with 7 of 81 (8.6%), is only 1.7-fold higher than the background (5.2%) (FET, p-value = 0.20). No comparable trend was found for CNVs having higher translation efficiencies.

### Inference of potential mechanisms of translational gene regulation

Previously published research has identified potential uORFs upstream of several genes that are amplified by CNVs in one or more of our strains, including *IRS4, ALY1, NTR2, KAE1, UIP5, TFA2, TGL4,* and *MLP1 (May et al. 2023)*. We developed *uorfish,* a uORF identification tool using a deep learning model to analyze ribosome profiling data from each strain. We identified 565 genes with high confidence uORFs genome wide, 216 of these exhibited significantly different translation efficiency in one or more strains (**Fig 3B**), which is significantly higher than expected at random (1.6-fold higher, HGM p-value < 1e-4). While genes amplified by CNVs have 1.3-fold higher uORF abundance than expected by chance this is not statistically significant (FET, p-value = 0.61), indicating that the amplification of genes is independent of the presence or absence of a uORF.

Recently, SSD1, an RNA binding protein, previously associated with aneuploidy tolerance (Hose et al. 2015; Hose et al. 2020) was identified as having targets specific to stress conditions using an analysis of high confidence binding sites in optimal and heat-shocked conditions (Bayne et al. 2022). To investigate the role SSD1 may play in gene regulation in evolved CNV strains we evaluated genes significantly enriched in the conserved ‘CNYUCNYU’ motif. Using the CNYUCNYU motif, we scanned the annotated TLs of the genome to identify genes enriched in the SSD1 motif, identifying 185 significantly enriched genes (FET, p-value < 0.05) of which 163 were present in our data set (Fig 4c).

**Fig 4.**
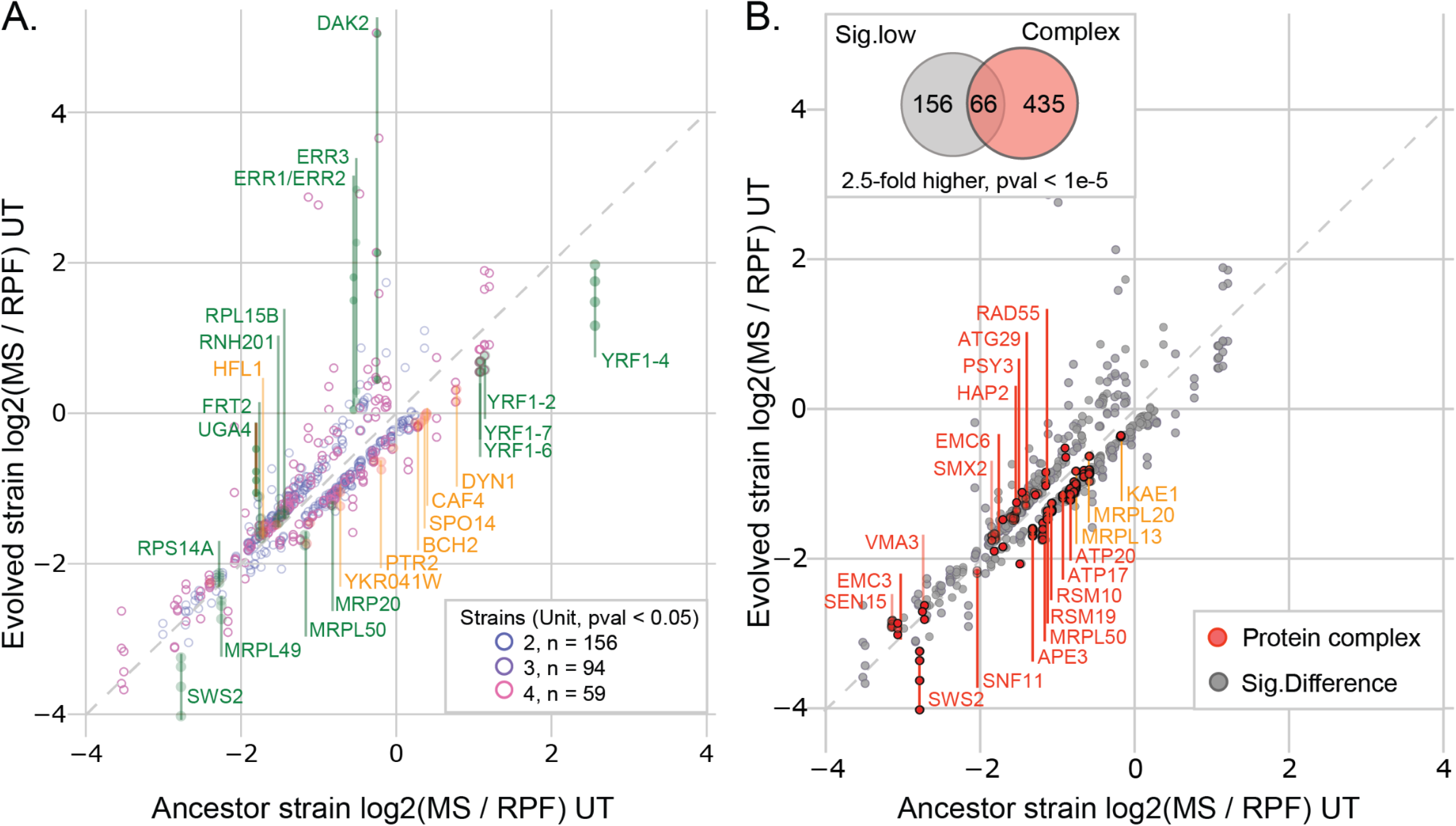
– Divergence in protein expression efficiencies and potential regulatory mechanisms. (**A**) 309 genes with significantly different protein expression efficiency relative to the ancestor in at least two evolved strains (unit transform test, p-value < 0.05). (**B**) Genes that have significantly different protein expression efficiency and are part of heterodimer protein complexes (red). Genes with significantly lower protein expression efficiency are 2.5-fold higher than random, HGM, p-value < 1e-5).

We found genes with uORFs were also enriched in SSD1 binding motifs in their TLs (2.4-fold higher, HGM, p-value < 1e-4). We also found that genes with significantly different translation efficiencies were enriched in SSD1 binding motifs (1.4-fold higher, HGM, p-value = 2.1e-3) and that genes with significantly different translation efficiency were 2.3-fold more likely to have both *SSD1* motifs and uORFs (HGM, p-value < 1e-5). Given the potential for interaction between uORFs and *SSD1* binding motifs in TLs, we used a relative genomic distance test co-occurrence (Favorov et al. 2012). We found that the proximity (relative distance of 0.0 – 0.1) of SSD1 motifs and uORFs is significantly higher than expected by chance (**Figure S4**).

One well known target of SSD1 is *UTH1*, a mitochondrial protein involved in regulating both mitochondrial biogenesis and degradation (Camougrand et al. 2004), the overexpression of *UTH1* has been shown to lead to cell death in yeast strains lacking SSD1 (Camougrand et al. 2003)) and is associated with the role of SSD1 in cellular response to aneuploid stress (Hose et al. 2020). Our analysis of the ribosome profiling data found a large abundance of uORFs and numerous CNYUCNYU motifs in *UTH1* (**Figure 3C)**. Similar co-occurrences of uORFs and CNYCNYU motifs were observed for several well described SSD1 targets including *SUN4*, *CTS1*, and *SCW10 (Bayne et al. 2022)*. While a manual analysis of ribosome profiling suggested that other well known targets of SSD1 such as *CLN2* (Ohyama et al. 2010)*, DSE2*, *NSA3*, *SRL1*, *SCW4*, *CCW12* (Bayne et al. 2022), and *TOS1* (Hose et al. 2020) these were not detected at the default quality threshold. Importantly, we also found that *SSD1* itself appears to have extensive uORFs and SSD1 motifs, suggesting it may auto-regulate (**Fig 3D**). Notably, *SSD1* has significantly lower mRNA abundance (p-value < 0.05) in every evolved strain compared to the ancestor; however, it is translationally upregulated in all evolved strains resulting in a final protein abundance slightly higher than the ancestor, consistent with translational regulation.

### Differences in protein expression efficiency

Differences in protein expression efficiency (i.e. the ratio of protein to ribosome abundance) can be evidence of post-translational regulation, likely the result of changes in protein stability (Senger et al. 2022; Ishikawa et al. 2020). However, unlike the translation efficiency estimation for which both RNA and RPF abundance measurements are produced using comparable methodologies and produce units of similar scale; protein abundance as measured by TMT-labeled mass spectrometry and ribosome profiling are substantially different making direct ratio calculations challenging. To address this challenge we first transform both MS and RPF abundances to a similar numerical scale while preserving features of the initial distribution, we then apply a statistical test of the distance between the protein expression efficiency in the evolved strain versus that in the ancestor (**Methods**). We define significance as any difference in expression that is greater than what is expected at random (for a similarly expressed gene) at a rate of 5% (eg. FDR 5%) or if the standardized residual of a linear regression is greater than 2 (**Table ST10**).

We find 379 (7.8%) of genes (**Fig 4A**) have significantly different protein expression efficiencies in evolved lineages compared to the ancestor, with 292 having significantly higher efficiency and 87 having lower efficiency. Of these 132 (35%) were significantly higher in multiple backgrounds, 32 (8.4%) were significantly lower in multiple backgrounds, and 29 (7.6%) were significantly different in opposite directions in specific strains.

We find 15 CNV amplified genes have significantly different protein expression efficiencies with the majority of these (12 or 80%) being significantly lower than the ancestor. Whereas only one CNV amplified gene, *HFL1*, is significantly higher in protein expression efficiency in multiple backgrounds, six CNV amplified genes; *YKR041W*, *PTR2*, *BCH2*, *SPO14*, *CAF4* and *DYN1*, are significantly lower in multiple backgrounds (**Fig 4B, orange**). Notably, although small in number, CNVs are 7.4-fold enriched in genes with significantly lower protein expression efficiency than the rest of the genome (FET, p-value = 5.8e-7), consistent with models of protein degradation as an important mechanism for dosage control (Ishikawa et al. 2017).

Previous studies of dosage compensation at the protein level have found evidence of targeted degradation of proteins involved in heteromeric complexes (Ishikawa et al. 2017), potentially through the ubiquitination of the exposed interfaces of unpaired heteromers (Ishikawa et al. 2020). To test this we looked for an enrichment of genes that encode proteins known to be involved in heterodimer complexes within the subset of genes with significantly lower protein expression efficiency. We found that genes with significantly lower protein expression efficiency were 2.5-fold enriched in proteins involved in heterodimer complexes (HGM, p-value < 0.05).

## Discussion

In this study we set out to characterize how gene expression changes at the transcriptional, translational, and protein level in response to adaptive CNVs that are selected in response to strong selection. We quantified different aspects of gene expression in 5 strains of yeast, 4 of which had undergone adaptation to glutamine-limited chemostats for hundreds of generations and had gained CNVs, as well as an ancestral strain unadapted to glutamine-limited chemostats and lacking any CNVs (**Fig 1**).

In a comparison of gene expression at the level of transcription and translation we observed over 2,000 genes had significant changes in expression and over 1,700 genes had significant changes in protein abundance (**Fig 2A**). The majority of genes amplified by CNVs had significantly higher levels of expression than their ancestral counterparts (**Fig 2B**) which agrees with previous reports on the broad effects of CNVs (Avecilla et al. 2023; Springer et al. 2010; Gasch et al. 2016). We also observed that the number of significantly divergent genes (including both amplified and unamplified) decreases at each successive level of expression, from a median of ∼1500 per strain for RNA, to ∼1100 for ribosome abundance, to ∼600 per strain for protein abundance. This agrees with previous reports of buffering of divergent gene expression at the translational (McManus et al. 2014; Kusnadi et al. 2022) and post-translational levels (Wang et al. 2018; Jiang et al. 2023). Furthermore, we find that many of these genes exhibit significantly different expression in multiple lineages (**Table ST4, ST5, ST6**), suggesting that the changes may be adaptive.

Since the level of expression at each stage of gene expression is dependent on the abundance of the molecule that precedes it, we sought to identify changes in gene regulation that alter the efficiency of expression at each level by normalizing each step of gene expression by the preceding step. This enabled us to evaluate changes in gene expression efficiencies between the evolved strains and the ancestor (**Fig 2C**, **Fig 3A, Fig4A**). By comparing the observed RNA abundance to what would be expected given the ancestral expression and the copy-number (**Fig 2C**) we found changes in transcription efficiency to be widespread with a median of 25% of all genes per strain. While CNVs had slightly higher rates of genes with either significantly higher or lower transcriptional efficiency, these were not significantly higher than the global background.

We found a median of 11% of all genes per strain exhibited significant changes in translation efficiencies relative to the ancestor. While considerably less than the 25% of genes with significant differences in transcription efficiencies, this decrease could either be due to the more limited role of changes in translational efficiency or a difference in the statistical powers of the two tests used.

Finally, we found that genes with significantly different protein expression efficiency are rare, with only a median 4% of all genes per strain. Given the conservative nature of the unit transform test this low percentage probably represents a lower-bound, with higher percentages, similar to those estimated previously being possible (Ishikawa et al. 2017). Although fewer genes exhibited lower protein expression efficiency, those that did were significantly enriched in genes amplified by CNVs in each strain, suggesting that decreased protein expression efficiency may be a general mechanism of dosage compensation for amplified genes. This potential role in mediating dosage compensation is in agreement with observations of natural isolates of yeast with adaptive aneuploidy (Muenzner et al. 2022) and human cancers (Schukken and Sheltzer 2022). One possible mechanism of dosage compensation at the protein level is the targeted degradation of stoichiometrically imbalanced proteins involved in heteromeric protein complexes (Ishikawa et al. 2020). Intriguingly, dosage sensitive genes that incur a fitness cost at high copy-number levels were also identified as being enriched in proteins involved in protein complexes (Makanae et al. 2013), suggesting potential limits to the capacity for this mechanism to compensate for dosage imbalances.

We found genes with significantly different translation efficiencies to be significantly enriched in upstream open reading frames (uORFs, **Fig 3B**) consistent with their established role in the translational regulation of gene expression (Ingolia et al. 2009; Chew et al. 2016; Moro et al. 2021). We found that genes within CNVs were not specifically enriched in uORFs. However, all genes with significantly different translation efficiencies were significantly enriched for uORFs, regardless of being CNV amplified or not. This suggests that global changes in gene expression in response to adaptation may be translationally mediated by uORFs but that this may not be a mechanism of dosage compensation specific to CNV amplified genes.

We also found that genes with significantly different translation efficiencies were also significantly enriched in binding motifs for the RNA binding protein SSD1 (**Fig 3B**) suggesting that it may also be involved in mediating changes in translation efficiency. Notably, *SSD1* has previously been shown to be a stress responsive RBP under glucose starvation (Bresson et al. 2020), heat shock (Bayne et al. 2022) and quiescence (Miles et al. 2019). It is capable of altering translation of bound RNA (Hogan et al. 2008; Ohyama et al. 2010) by preferentially binding to transcript leaders (TLs) and interfering with ribosome scanning (Bayne et al. 2022). Given this background, we explored the potential for interaction between uORFs and *SSD1*. Indeed, we find that genes with uORFs are significantly enriched in *SSD1* motifs in their TLs and that these motifs have a closer proximity to the uORFs than expected by chance (**Figure S4**). These findings suggest that SSD1 may act as a stress responsive *trans*-acting factor driving uORF mediated translational regulation, potentially similar to the sex-lethal (SXL) protein binding that controls uORF translation of *msn2* in *Drosophila melanogaster* (Medenbach et al. 2011).

Our findings help contextualize the role of SSD1 in CNV and aneuploid yeast strains. Previous studies have shown a much higher level of gene specific dosage compensation in aneuploid yeast strains with a functional copy of *SSD1* (such as the S288c strain, as was used in this study) compared to yeast strains lacking a functioning *SSD1* (such as W303). This suggests that the importance of SSD1 in mitigating the fitness defects of CNVs may be a consequence of the broader role SSD1 plays in the translational regulation of gene expression under stress, and not specific to CNVs per se. Indeed, this may explain some of the observed but varying and incomplete overlap seen in SSD1 targets, genes involved in the hypo-osmotic stress pathway (Bayne et al. 2022), CAGE (Tsai et al. 2019), and analyses that sought to identify the CNV and aneuploid stress response at the transcriptional level (Avecilla et al. 2023).

Our study is not without its limitations. Notably, the number and type of strains evaluated limit the scope of our inferences and the generalizability of our results. Beyond that, technical limitations, such as ratio compression of TMT-labeled mass spectrometry data (Ting et al. 2011) and statistical tests of changes in expression across multiple levels make certain hypotheses hard to test. However, this work points to several avenues for ongoing research. While we have shown that gene expression, at the transcriptional and post-transcriptional levels, is significantly different from the ancestor in glutamine-adapted strains, it is unclear how rapidly these changes arise, how recurrent they may be, and how they change over the course of long term evolution. Furthermore, the potential for RBPs to interact with uORFs, and thereby alter downstream gene expression, while not unprecedented (Medenbach et al. 2011; Spealman et al. 2018) largely unexplored.

## Materials and Methods

### Strains and growth media

Strains evaluated in this paper were originally published in (Lauer et al. 2018), they include both long-term experimentally evolved strains as well as their ancestor. The naming convention is as follows: Ancestor (DGY1657); Trans (DGY1726); ODIRA_A (DGY1735); ComQuad (DGY1741); and ODIRA_B (DGY1743). Trans and ODIRA_A are from population samples after ∼150 generations, ComQuad and ODIRA_B are from population samples after ∼250 generations. All strains are derived from FY4 (BY1747, **Figure S1A**) and modified by the inclusion of a GFP and KanMX reporter cassette (**Figure S1B)**. All evolved strains were evolved in miniature chemostats under minimal glutamine growth media for either 150 or 250 generations, as described previously. For this study, strains were struck out on YEPD plates from –80°C freezer stocks and grown for 2 days at 30°C. Single colonies were selected based on the presence of fluorescence produced from the previously described mCitrine reporter (Lauer et al. 2018).

### Illumina genomic DNA sequencing

For short-read Illumina sequencing of strains we relied on previously generated data ((Lauer et al. 2018), NCBI SRA accession SRP142330). Briefly, we performed genomic DNA extraction (Hoffman and Winston 1987), followed by quantification using SYBR Green I and standardized to 2.5 ng/μL. Libraries were constructed using Illumina Nextera tagmentation (Baym et al. 2015). Final concentrations were measured using SYBR Green I, fragment sizes were measured using Agilent TapeStation 2200 before being balanced and pooled. Illumina DNA libraries were sequenced on an Illumina NextSeq 500 using 2×75 paired-end protocol.

### Illumina genomic DNA alignment and genotyping

All sequences were aligned against the Ensembl_R64.1.50 reference genome. We aligned reads using bwa mem (v0.7.17, (Li and Durbin 2009)) and generated BAM files using samtools (v1.14, (Danecek et al. 2021)). FASTQ files for all sequencing are available from the SRA (accession SRP142330).

We used GATK4 HaplotypeCaller (ver 4.1.9.0, (Van der Auwera et al. 2013)) to identify SNPs and indels in 4 evolved strains and their ancestor (DGY1657). HaplotypeCaller was run using a haploid ploidy with default settings against the Ensembl S. cerevisiae R64-1-1 reference genome. Variants were filtered using GATK VariantFiltration: SNPs (QD < 2.0, FS > 60.0, MQ < 30.0, SOR > 4.0, MQRankSum < –12.5, ReadPosRankSum < –8.0); Indels (QD < 2.0, FS > 200.0, SOR > 10.0); (**Table ST11**). Variants were further filtered using Herança (v.0.8; https://github.com/pspealman/heranca), a lineage aware quality control script that filters variants more likely explained as sequencing errors (**Figure SF2**). Variants were annotated and predicted effects scored using Ensembl VEP (release 107, (McLaren et al. 2016) default settings, *S. cerevisiae* R64-1-1, Upstream/Downstream distance (bp): 500). Potential structural and copy-number variants using Illumina paired end data were identified using our custom analysis tool CVish (v1.1, (Lauer et al. 2018)). **Table ST12**)

Potential SNV and indel identification of each strain was performed using GATK’s HaplotypeCaller (v4.1.9.0, (Van der Auwera et al. 2013)) in single-sample mode and annotated using Ensembl VEP (release 107, (McLaren et al. 2016)), These are reported in the supplement (**Table ST11)**. Potential structural and copy-number variants using Illumina paired end data were identified using our custom analysis tool CVish (v1.1, (Lauer et al. 2018)). **Table ST12**)

### Nanopore genomic DNA sequencing, alignment, and genotyping

All yeast strains were grown to greater than 1 × 10^7 cells/mL in 300 mL Glutamine minimal media. Genomic DNA from each strain was extracted using Qiagen 20/G Genomic tips from ∼1.5 × 10^9 cells using the manufacturer’s protocol.

All genomic DNA was barcoded using Oxford-Nanopore’s native barcoding genomic DNA kit (EXP-NBD104), adapters were added using the ligation sequencing kit (SQK-LSK109). The manufacturer’s protocol (versions NBE_9065_v109_revB_23May2018 and NBE_9065_V109_revP_14Aug2019) was followed with the following exceptions: incubation times for enzymatic repair step were increased to 15 min. All Agencourt AMPure XP beads were incubated for 30 min at 37°C before elution. Adapter ligation time was increased to 10 min. Multiplexed libraries were loaded on MinION flowcells (FLO-MIN106D R9) and run on a MinION sequencer (MIN-101B). (Quinlan and Hall 2010)

All sequences were aligned against the Ensembl_R64.1.50 reference genome (Cunningham et al. 2022). ONT long-read sequences were aligned using minimap2 (v2.22, (Li 2018)), with variant detection using sniffles2 (v2.0.6, (Li 2018; Sedlazeck et al. 2018)), potential ODIRA CNV breakpoints were evaluated with mugio (v1.7, (Spealman et al. 2020)). DNA depth across genomes was calculated per nucleotide using bedtools ‘genomecov’ (ver 2.29.2, (Quinlan and Hall 2010)). These were used in conjunction with the CNV breakpoints identified by CVish to manually reconcile the CNV topologies for each strain (**Fig S1, Table ST1).**

### DNA reads and depth estimation

We identified ∼15.4% of genes in the ancestral strain had long-read DNA read-depth not near one copy. Some of these are known deletions (*MATALPHA*, *SRD1*) or are regions with known properties of known copy number expansion (*CUP1* region, ENA region, and ASP region) or transposons. To prevent the over or underestimation of expected expression in evolved strains, we manually analyzed the locus and flanking regions for signs of CNV breakpoints using both short and long-reads. If no evidence was found for a breakpoint the locus was assigned a copy-number of 1 (**Table ST1)**.

### Growth conditions and cell harvesting

Colonies were grown overnight on YEPD then used to inoculate 500 mL of minimal glutamine growth media in chemostats. These were then grown to saturation (approximately 48 hours ∼1e7 cells / mL, measured using Coulter Counter) at 30°C in aerobic conditions. After saturation was achieved chemostats were switched to continuous mode with an inflow rate of fresh minimal glutamine growth media at a rate of 0.12 hour / L (corresponding to a population doubling time of ∼5.8 hours). This was maintained for 24 hours before the cells were harvested. This was performed in duplicate for each strain for the simultaneous extraction of RNA and RPFs. This was performed for 5 replicates for the generation of material for mass spectrometry.

### RNA-seq and ribosome profiling

Cells were harvested following previously described methods (Spealman et al. 2016). Briefly, cycloheximide was added to the media to a final concentration of 100μg/L and incubated for 2 min, before being separated from the media using rapid vacuum filtration using Millipore 0.22μm filters. Cells were resuspended from filters using polysome lysis buffer with cycloheximide (CHX) and then flash frozen immediately in liquid nitrogen before being transferred and stored at –80°C. RNAseq and ribosome profiling was performed by TB-SEQ, Inc. (Palo Alto, CA). As per (McGlincy and Ingolia 2017), cells were lysed in polysome lysis buffer with CHX, cells for each sample were split into RNA and RPF aliquots. For RPF aliquots, RNase I was used to digest polysomes, monosomes were purified using a sucrose cushion. Fragment size selection of 25-32 nucleotides was performed using a polyacrylamide gel. No ribosomal depletion or poly-A tail selection was made to either RPF or RNA libraries. Fragment size and concentrations were measured using bioanalyzer and qubit before sequencing using Illumina NovaSeq6000, single read, 1×50 cycles.

### Mass spectrometry and analysis

For the generation of material for MS 5 biological replicates for each strain were grown in chemostats as described above. Cells were collected from the chemostats using the same techniques described above. TMT-labeling and LC-MS were performed by Proteomics Laboratory, NYU Langone Medical Center (New York, NY). We used 16Plex TMT labeling to enable pooling of peptides from 16 samples (Ch1-16) for offline fractionation and LC-MS/MS detection and quantification (Figure S. Five replicates were generated for each strain and the total run was performed in two batches (“A”,“B”). Ch1 was used as the common reference standard, an equimolar mix of all the samples, labeled with TMT Ch1. All samples were lysed using a TFA approach (Doellinger et al. 2020). Both batches were fractionated by offline HPLC chromatography using reverse phase C18 stationary phase at high pH. Peptides from collected fractions were separated by online HPLC on C18 at low pH coupled to the MS instrument.

### Gene expression analysis

RNA and RPFs were aligned using STAR (ver 2.7.6a, (Dobin et al. 2013)) filtering for known non-polyadenylated ncRNA (snRNA, rRNA, and tRNA) using Ensembl_R64.1.1 ncRNA fasta (**File SF1**). Counts per gene were calculated using bedtools (v2.29.2, (Quinlan and Hall 2010)) coverage –counts –s –b against Saccharomyces_cerevisiae.R64-1-1.50.gff3. For mass spectrometry the spectra were analyzed using MaxQuant (v2.1.0.0, (Tyanova, Temu, and Cox 2016)) with parameters described previously (Yu et al. 2020). Parameter and configuration files are provided as supplementals (**File SF2, SF3**).

Spearman *rho* correlations between replicates for each level of expression for each strain were calculated (**Table ST2**), for RNA median rho between replicates (*rho* = 0.9878), RPF (*rho* = 0.9879), and MS (*rho* = 0.9965), which is sufficiently high for replicates. Between levels (but within the same strain) we found the median rho for RNA to RPF (*rho* = 0.6271), RPF to MS (*rho* = 0.6012), RNA to MS (*rho* = 0.5453). We also compared between strains (**Table ST3**) and found the strains to be very well correlated with the median rho for RNA (rho = 0.9396), RPF (rho = 0.9258), and MS (rho = 0.9915).

Differential gene expression analysis for RNA and RPF abundances were calculated DESeq2 (v1.6.3, (Love et al. 2014)). Results of DESeq2 are available as supplemental files (**RNA, Table ST4; RPF Table ST5**). Significance is defined as Benjamini-Hochberg (BH) adjusted p-value < 0.05.

Differential gene expression analysis for mass spectrometry data was performed using Perseus (v1.6.15.0, (Tyanova, Temu, Sinitcyn, et al. 2016)). Data preprocessing and normalization procedures were performed as described (Yu et al. 2020). The statistical test for significance in differential abundance is Welch’s t-test of the difference between the normalized protein group abundances in each strain relative to the ancestor. Significance here is defined as q-value < 0.05 using BH multiple hypothesis test correction (ie. BH adj.p-value < 0.05), (**Table ST6**). Volcano plots indicating the protein abundances of CNV in each strain are in the Supplemental (**Figure SF5, SF6, SF7, SF8**).

Gene ontology term (GO term) enrichment was calculated using Yeastmine (Balakrishnan et al. 2012) with significance cutoff at BH adj-p-value < 0.05. Redundant terms were then reduced using REVIGO (Supek et al. 2011) using the Tiny option, UNIPROT *Saccharomyces cerevisiae* S288c database, and SimRel semantic similarity measure. Heterodimer protein complexes were compiled from Yeastmine (Balakrishnan et al. 2012) with the criteria that were categorized as complexes and had annotated stoichiometry values, these were combined with a manually curated list from (Taggart and Li 2018), and with complexes specified by (Ishikawa et al. 2017).

All Venn diagrams were originally visualized using Venny (v2.1.0, https://bioinfogp.cnb.csic.es/tools/venny/), proportional Venn diagrams were generated using DeepVenn (Hulsen 2022). All expression track data was visualized using IGV browser (v2.9.4, (Robinson et al. 2011)).

For quality control purposes we set an arbitrary expression threshold. Genes were required to not be transposon element associated genes, have at least 100 aligned RNA reads, 10 RPFs, and have a significant majority peptide fragment detected by MS. This reduced our gene set to 4291 genes in total.

### Transcription efficiencies calculated as observed versus expected

Transcription efficiency was calculated using evolved to ancestor strain pairwise tests of observed to expected RNA abundance using DESeq2, using two replicates each. Expected RNA abundances were calculated using a custom script that multiplies the observed abundance in ancestor by the copy-number of that gene in the evolved strain. Differential abundance analysis was performed on both unnormalized observed and unnormalized expected RNAseq reads using DESeq2 (v1.6.3, (Love et al. 2014)). Results of DESeq2 are available as a supplemental file (**Table ST8**). Significance here is defined as BH adj.p-value < 0.05 as calculated by DESeq2.

### Translation efficiencies calculated using ratiometrics and FET

Translation efficiency was calculated using evolved strain to ancestor strain pairwise tests of RPF/RNA ratios. RPF and RNA abundances are first normalized using TPM (**Figure SF5**). Replicates were combined by taking the median RPF and median RNA separately before assessment as a ratio. Evolved RPF/RNA ratios per gene were compared to the ancestral RPF/RNA ratios using Fisher’s Exact Test (FET). Results of FET evaluation of translation efficiency are available as a supplemental file (**Table ST9**).

### Unit transform

Ideally, protein expression efficiency would be calculated using a ratiometric method similar to translation efficiency. However, while ratiometric analysis is robust for similarly scaled data it may generate spurious results for data with excessively different scales. Because ribosome profiling reports numbers 2 to 3 magnitudes higher than those reported by mass spectrometry we need to first transform the measurements to similar scales before proceeding.

While there are many approaches for data transformation, we opted to use Box-Cox power transform (**Figure SF5**) as implemented in scikit-learn (Pedregosa et al. 2011) as it retained meaningful features of the original distribution. We next transformed the data by MinMax scaling to a range of (0,1] (Pedregosa et al. 2011) so that all values greater than zero.

### Unit transform test

To determine if differences in unit transformed expression data are significant we require the data meets one criteria and we apply two orthogonal tests. First, we require that the variance in expression between replicates can not exceed the distance between strains. Secondly, after fitting a linear least-squares regression (lm function (R Core Team 2019)) of the evolved and ancestor expression, statistically significant outliers are then we then identified as those with standardized residuals (rstandard function (R Core Team 2019)) greater than 2. To complement this we also perform a second analysis.

First, we rank all genes using the difference of their expression measurements (**Figure SF6**). Then, we score each gene by taking the ratio of their expression. To determine if this score is greater than expected by chance we randomly sample the nearest 10% of genes of similar rank and count the number of times the score is met or exceeded by similarly ranked genes. A reported FDR of 0.05 means that 5% of the genes out of a random sampling of similarly ranked genes, in a simulation conducted 1000 times, met or exceeded the score being evaluated.

One way to evaluate the unit transform test method is to benchmark it against results from previous tests. In a performance comparison to DESeq2 data using unnormalized RNA abundances we find that the unit transform test performs very well (**Figure SF6, SF7**) with a false positive rate of 0.01, a false negative rate of 0.35 and a F1 score of 0.73. Performance decreases slightly when compared to the application of FET to TPM normalized translation efficiency ratios, where the unit transform test has a false positive rate of 0.03, a false negative rate of 0.34 and a F1 score of 0.63 (**Figure SF8, SF9**). In both cases the False Positive rate is appreciably low, suggesting the unit transform test method is sacrificing sensitivity for the precision.

### Protein expression efficiency calculated using unit transform and test

For each evolved strain we compared the protein expression efficiency ratio (Unit transformed MS intensity / Unit transformed RPF abundance) to the Ancestor ratio (Figure SF5). Replicates were combined by taking the median MS and median RPF separately before assessment as a ratio. Significant difference between the evolved and ancestor strains was calculated using the unit transform test at an of FDR < 0.05. Results of the unit transform test evaluation of translation efficiency are available as a supplemental file (**Table ST10**)

### Identification of high confidence predicted uORFs

Because uORF activity is known to be condition sensitive (Hinnebusch 2005) (Moro et al. 2021) and no previous study has been conducted using glutamine-limited growth media or CNVs, we sought to identify them using our ribosome profiling data. To identify ribosome occupied uORFs in these datasets we developed uORFish (github.com/pspealman/uorfish), a deep neural network uORF predictor trained on either ‘validated’ or ‘predicted’ uORFs from (Spealman et al. 2018) or ‘validated’ uORFs from (May et al. 2023). Note that the validation methodology used in (May et al. 2023) was unable to assess uORFs greater than 180bp distance from the main ORF TIS.

To prevent training on uORFs that are not active under the conditions used, uORFs are required to have ribosome occupation inframe to potential alternative translation initiation sites. All ribosomes were assigned to reading frames using p-site fractionation (Spealman et al. 2018). These viable training uORFs were used to train a DNN using pytorch, (Paszke et al. 2019) for uORF identification for each ribosome profiling replicate. High confidence predicted uORFs were defined as uORFs with prediction scores greater than 0.9 present in both replicates. Predicted scores for all candidate uORFs with scores greater than 0.5 are available as supplemental files (**Table ST13**).

### SSD1 motif analysis

To identify genes enriched in SSD1 motifs in their TLs we first generated a list of TLs for all genes using the Yeastmine (Balakrishnan et al. 2012) annotated ‘Five Prime UTR’ (Pelechano et al. 2013) or from (Spealman et al. 2018), taking the longest recorded instance. Combined these encompassed 6119 genes, 6080 of which exceeded the length of the motif. We converted these to fasta using bedtools ‘getfasta’ (ver2.29.2, (Quinlan and Hall 2010)). These were then scanned for perfect matches to the CNYUCNYU motif (Bayne et al. 2022), irrespective of strand. The background rate, genome wide was calculated per nucleotide as 2.39e-3, the background rate using only TLs was 2.91e-3 per nucleotide. Significance was calculated using FET, using the proportional ratio of (observed hits of TL / length of TL) versus (Total hits / Total length), (**Table ST14**). We found that of the 219 genes that were significantly enriched in SSD1 motifs, 163 were present in our data set.

### Data and resource availability

All data is publicly available. Illumina short-reads are available on NCBI SRA (PRJNA451489). ONT Nanopore long-reads are available on NCBI SRA (PRJNA591579). RNA-seq (GSE246093) and ribosome profiling (GSE246094) data are available through NCBI GEO. Mass spectrometry data is available through PRIDE (PXD046587).

All scripts and data used in the analysis included in this paper are available via GitHub: https://github.com/GreshamLab/adaptive_gene_expression

## Acknowledgements

We are grateful for funding from NIGMS (R35GM153419, R01GM134066), NSF (1818234), and BSF (2021276) (DG) and NIGMS (F32GM131573) (PS). We thank members of the Gresham lab for their insight and feedback.

