## Supplemental_Materials for "Multilevel gene expression changes in lineages containing adaptive copy number variants"

### Supplemental figures:

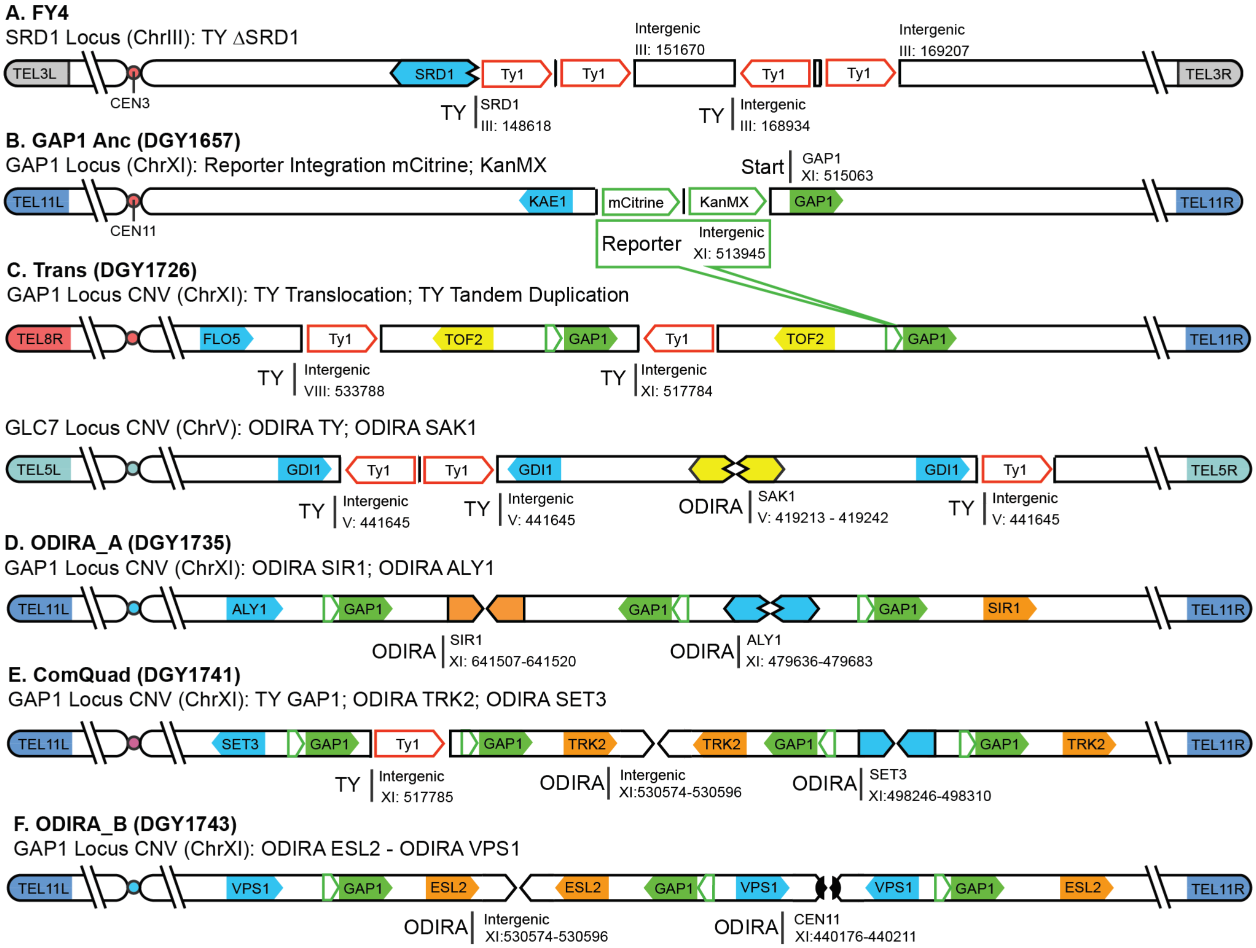


**Figure S1. Schematic of CNV Topology**

(**A**). Diagram showing the disruption of SRD1 in FY4 (**B**). Diagram showing the integration of the reporter in DGY1657 (**C**), the reporter is represented by a green box in all subsequent diagrams. Topology diagrams for evolved strains indicating CNV breakpoints, orientations, and the occurrence of transposon events **C-F**). CNV breakpoints are annotated with their most likely mechanism: transposon-yeast (red arrow), origin-dependent inverted-repeat amplification (ODIRA). Gene copy-number values for each strain are available in Supplementary Table 1.

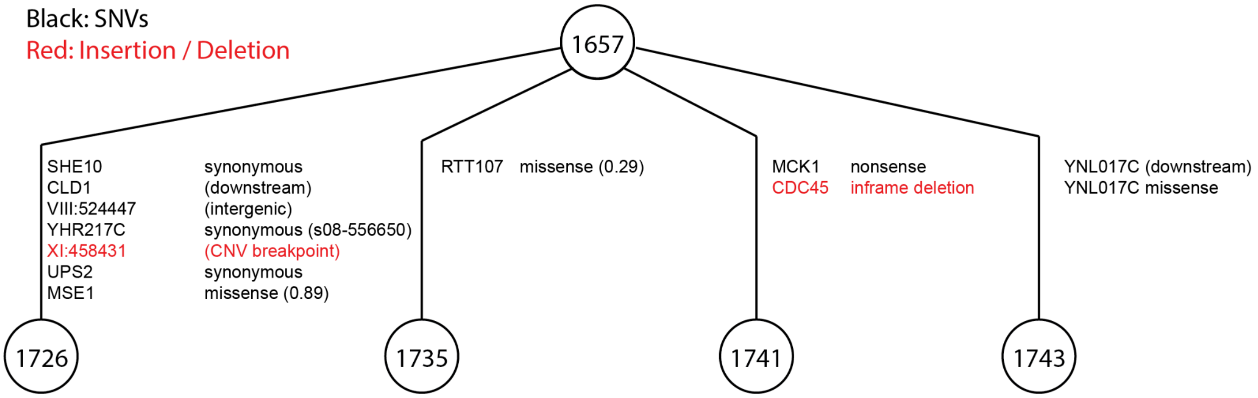


**Figure S2: Mutations in evolved strains**

Schematic representing the variants in the evolved strains versus their ancestor. Each variant in black is a SNP while variants in red are Indels. Variants within the CDS of a gene are marked as synonymous, nonsense, or missense. Missense variants also have an estimated mutational effect value (SIFT) when possible with lower values being more severe. In only one case was a CNV breakpoint identified using GATK (XI:458431, DGY1726). All variants are novel (ie. not in Ensembl Variant Catalog) except for the synonymous mutation of YHR217C in DGY1726

**Single nucleotide variants and indels analysis**

Potential SNV and indel identification of each strain was performed using GATK's HaplotypeCaller (v4.1.9.0, (Van der Auwera et al. 2013)) in single-sample mode and annotated using Ensembl VEP (release 107, (McLaren et al. 2016)), (Table ST11). Variants were further filtered using Herança (v.0.8, [2]), a custom lineage aware quality control script that filters variants more likely explained as sequencing errors These are reported in the supplement (**Table ST11)**.

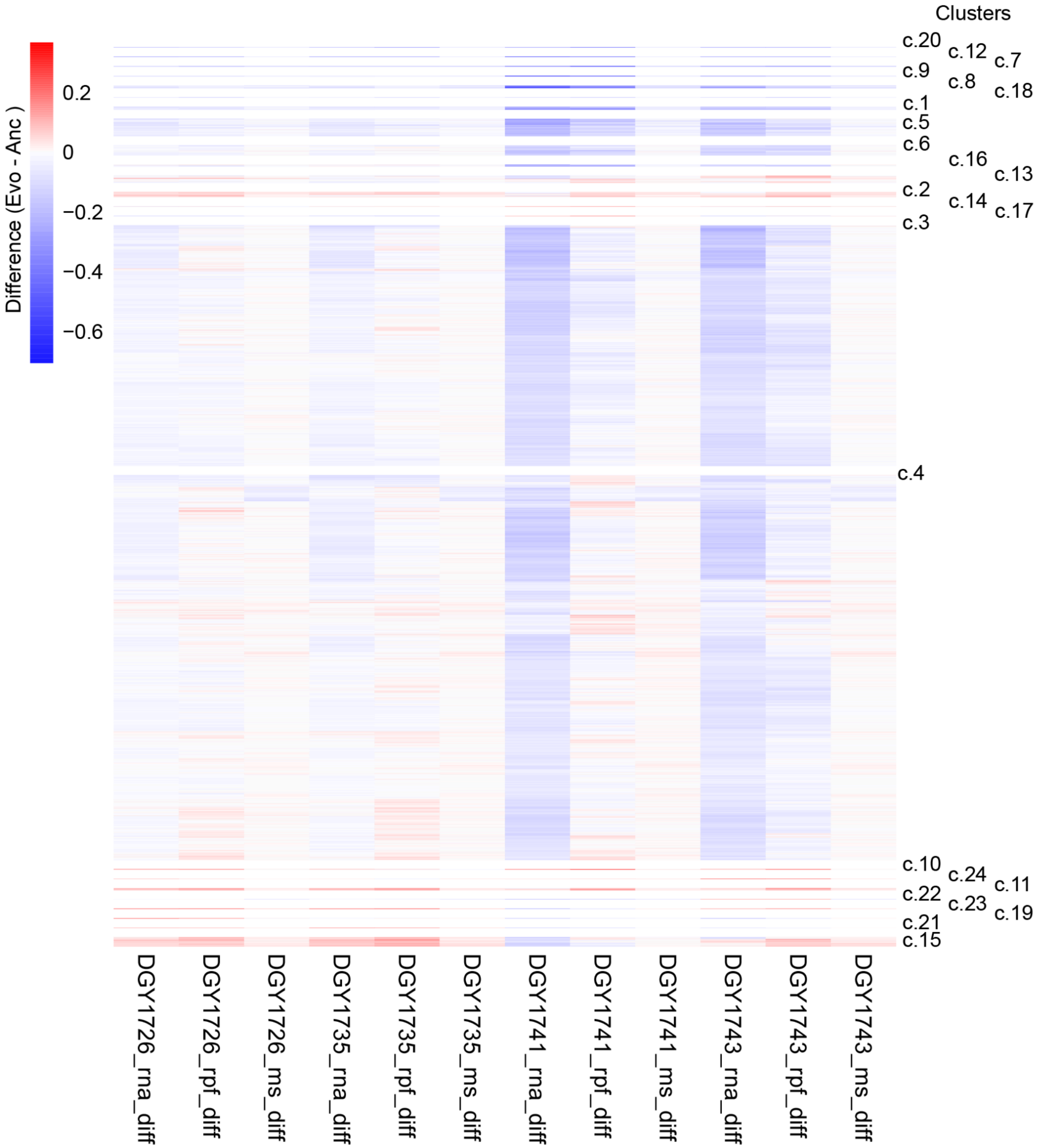


**Figure S3: Heatmap of multi-level expression with clustering (k=24).** Heatmap shows the difference (Evolved - Ancestor) of the Unit Transformed expression data for mRNA, RPF, and MS intensity for all evolved strains.

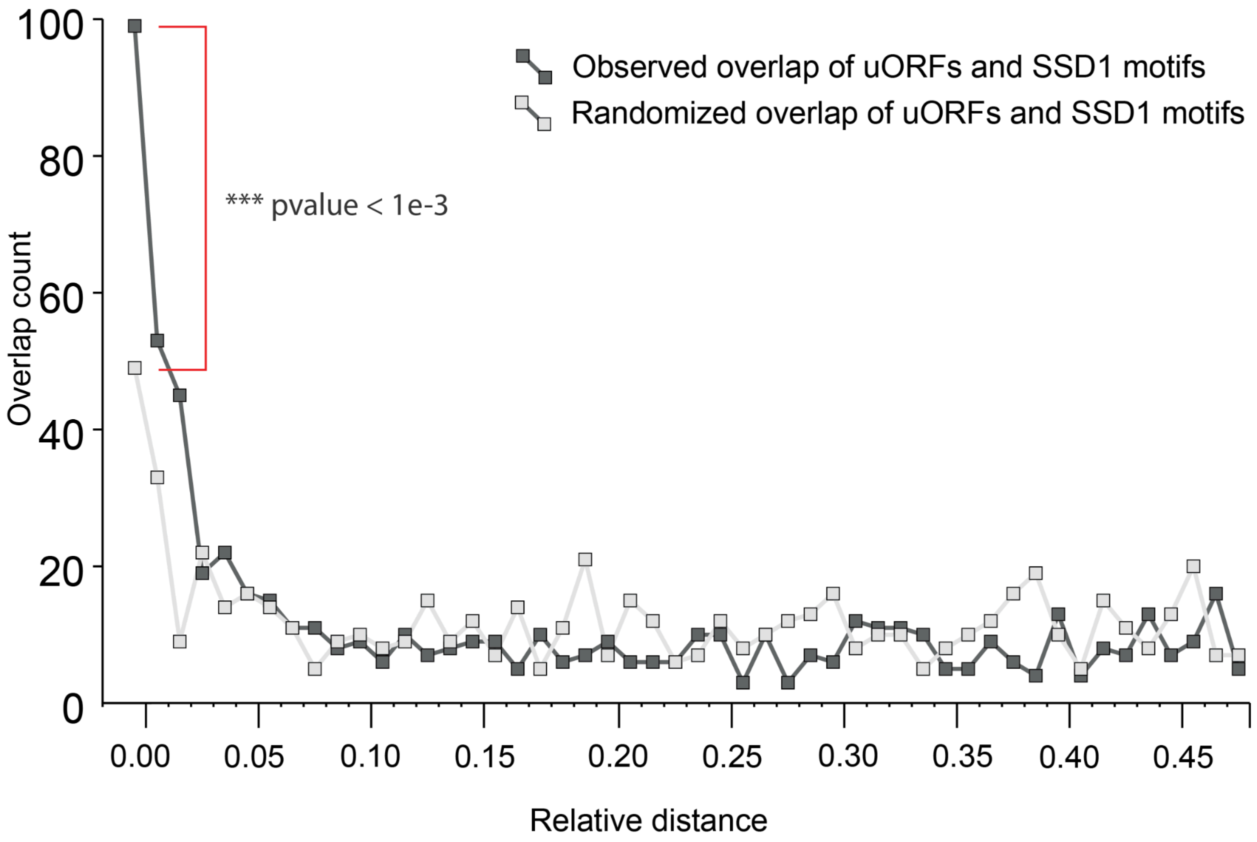


**Figure S4: Relative distance plot:** Showing the relative distance (Favorov et al. 2012), between overlapping uORFs and SSD1 motifs in transcript leaders. The observed (dark gray line) overlap in the immediate proximity (relative distance 0.00) is significantly higher than expected by chance (light gray line). Calculated using bedtool's reldist (Quinlan and Hall 2010).

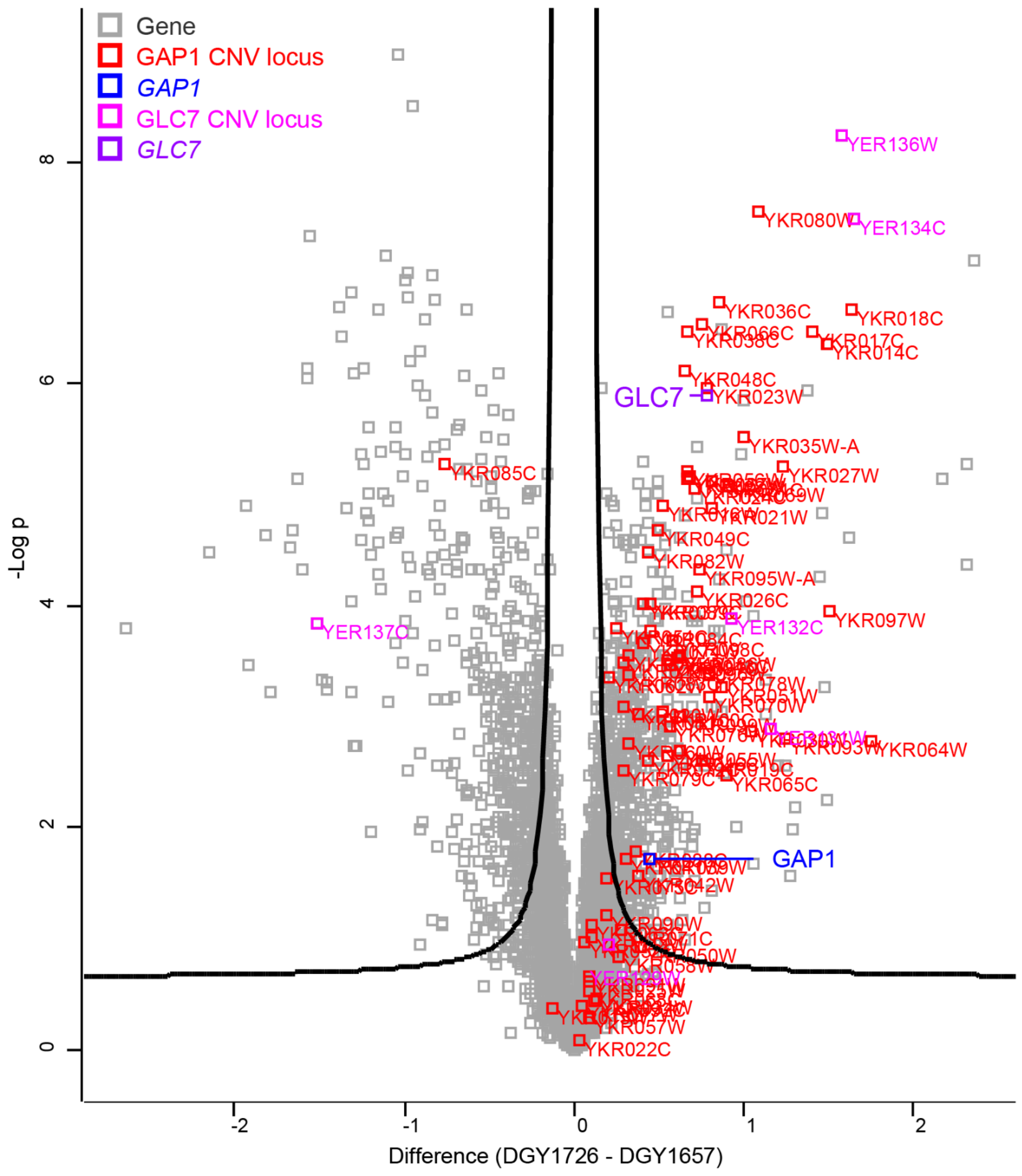


**Figure S5: Differential Protein Abundance Results for Trans (DGY1726).** Volcano plot of results of Welch's t-test between Trans (DGY1726) and Ancestor (DGY1657). Two regions of the genome have been CNV amplified. Genes amplified with the GAP1 locus are shown in red, *GAP1* is shown in blue. Genes amplified with the GLC7 locus are shown in pink, *GLC7* is shown in purple.

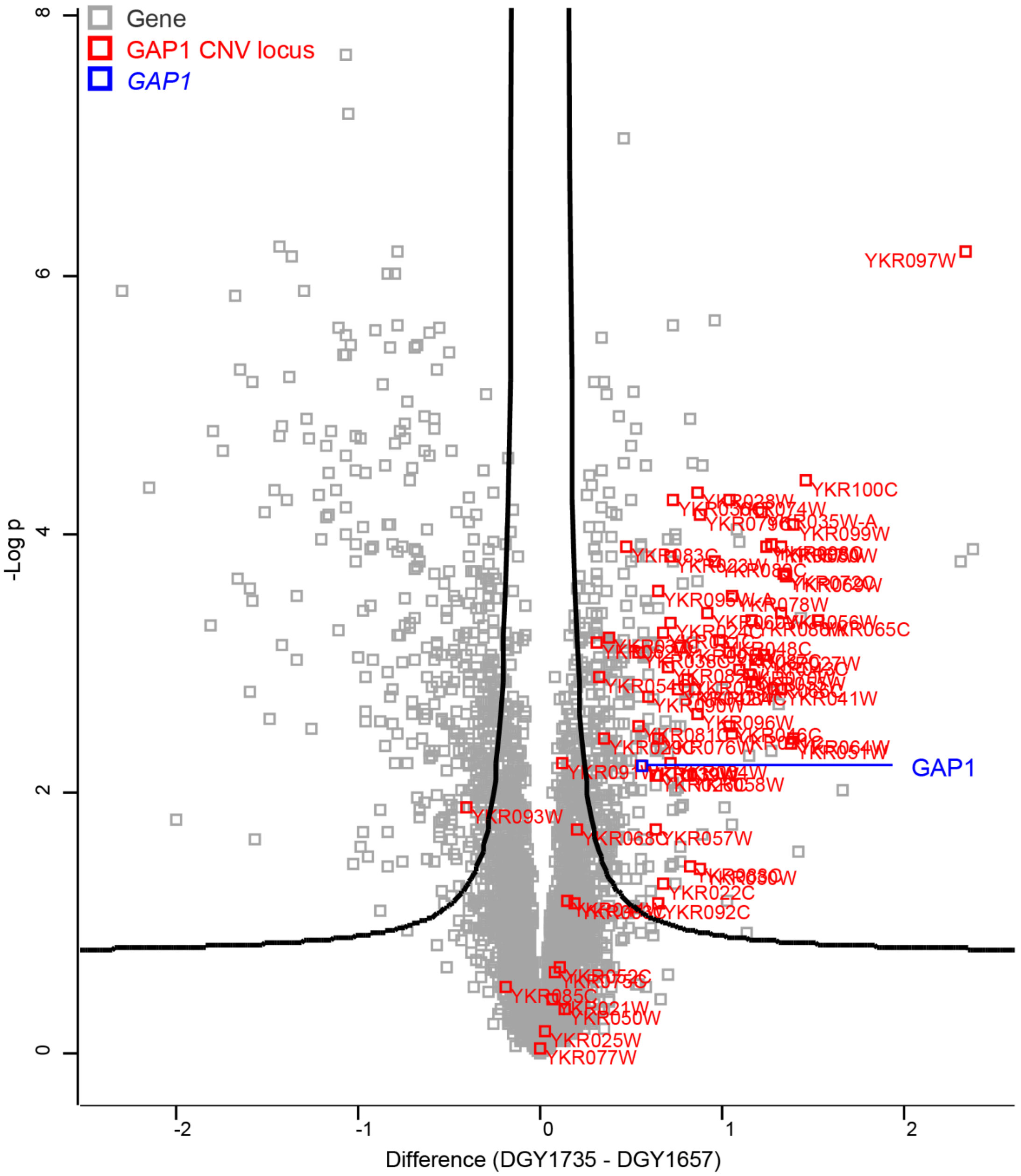


**Figure S6: Differential Protein Abundance Results for ODIRA_A (DGY1735).** Volcano plot of results of Welch's t-test between ODIRA_A (DGY1735) and Ancestor (DGY1657). Genes amplified in the GAP1 locus are shown in red, *GAP1* is shown in blue.

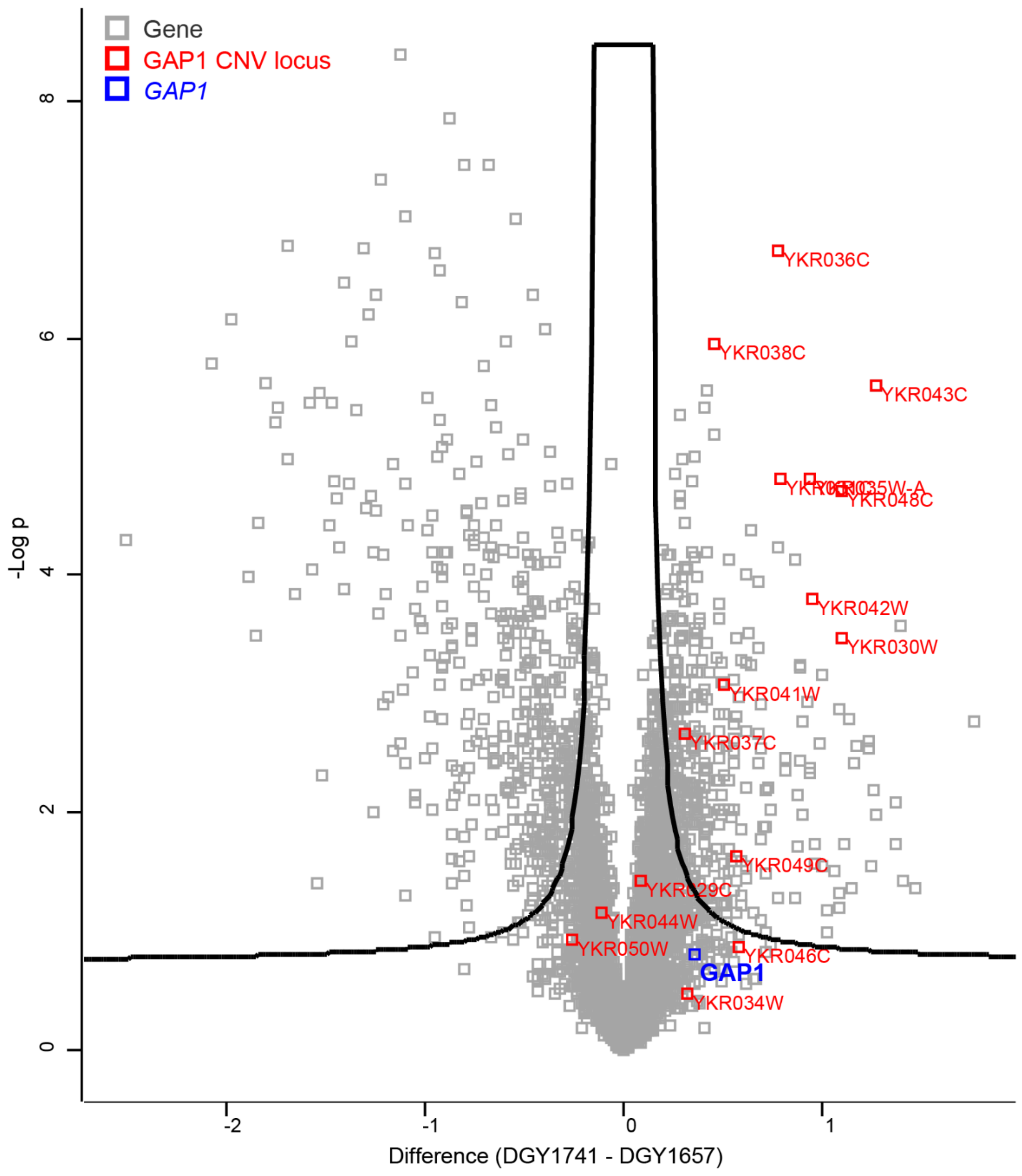


**Figure S7: Differential Protein Abundance Results for ComQuad (DGY1741).** Volcano plot of results of Welch's t-test between ComQuad (DGY1741) and Ancestor (DGY1657). Genes amplified in the GAP1 locus are shown in red, *GAP1* is shown in blue. Note that *GAP1* protein abundance in ComQuad is not significantly different from the Ancestor despite having been amplified to a copy-number of 4.

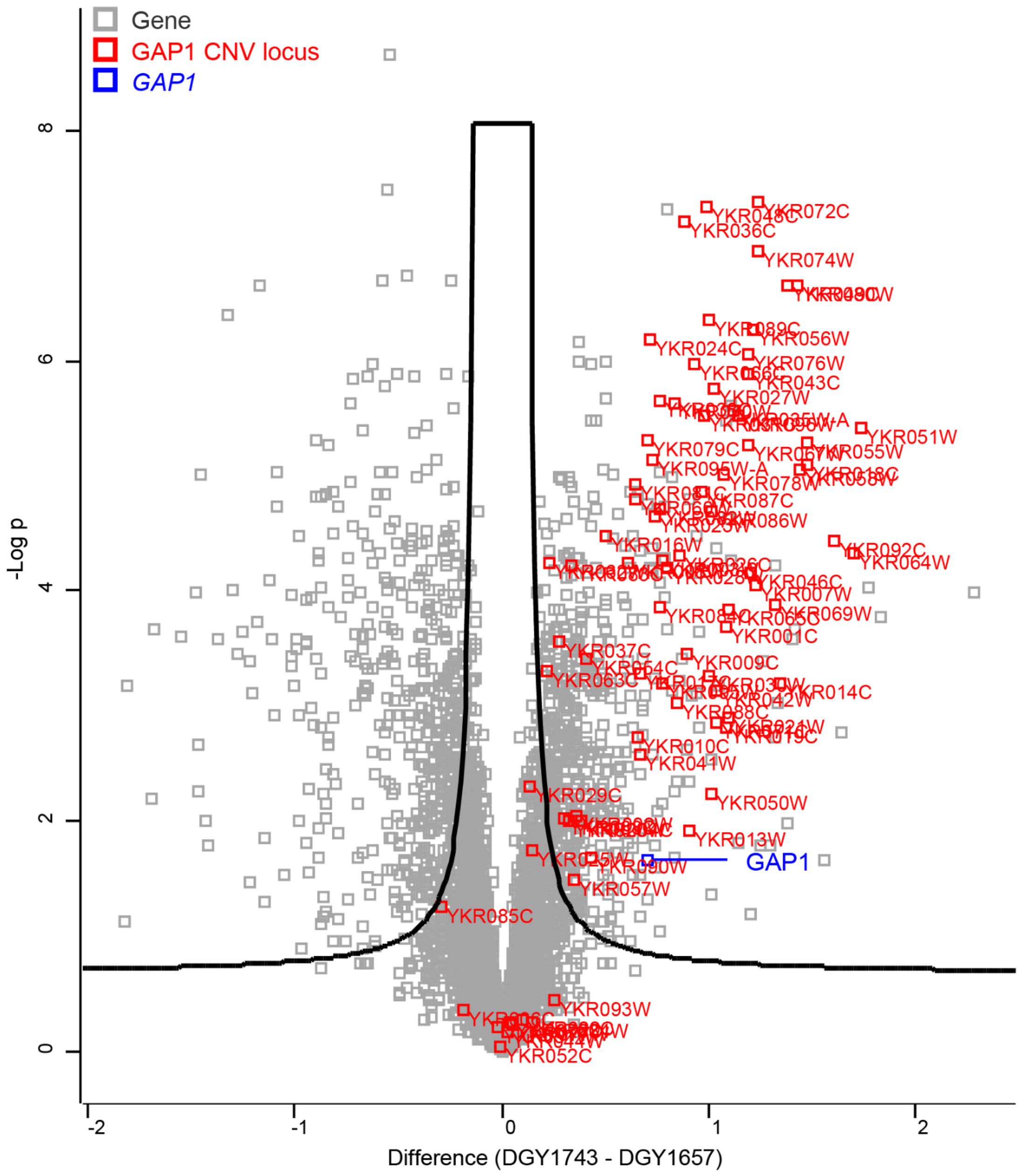


**Figure S8: Differential Protein Abundance Results for ODIRA_B (DGY1743).** Volcano plot of results of Welch's t-test between ODIRA_B (DGY1743) and Ancestor (DGY1657). Genes amplified in the GAP1 locus are shown in red, *GAP1* is shown in blue.

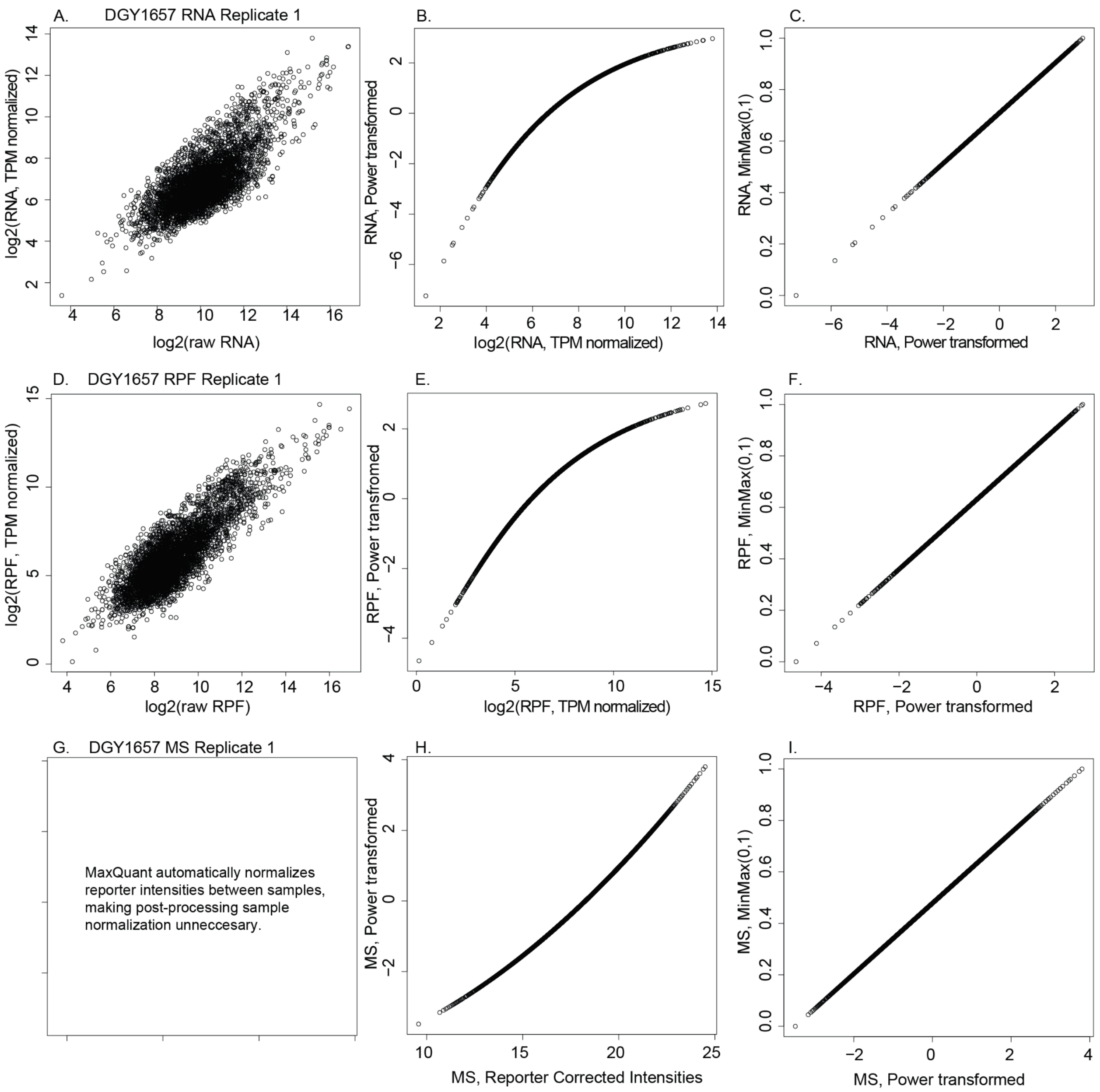


**Figure S5: Example of Power Transform of Expression Data.** Large numerical differences in scale separate expression data, with RNA and RPF ranging from 10-100,000 read counts per gene compared to 10-25 peptide reporter intensities from MS data. To compare these directly we sought to transform the data into similarly scaled units. For RNA and RPF data (A,D) we first normalize between samples for sequencing depth using TPM (10.1186/1471-2105-12-323). Next (B,E,H), we bring measurements within a similar scale using a Box-Cox Power Transform (http://www.jstor.org/stable/2984418), followed by a MinMax transformation (C,F, I) to keep all values positive. Both Powertransform and MinMax were implemented using Scikit-learn (https://jmlr.csail.mit.edu/papers/v12/pedregosa11a.html)

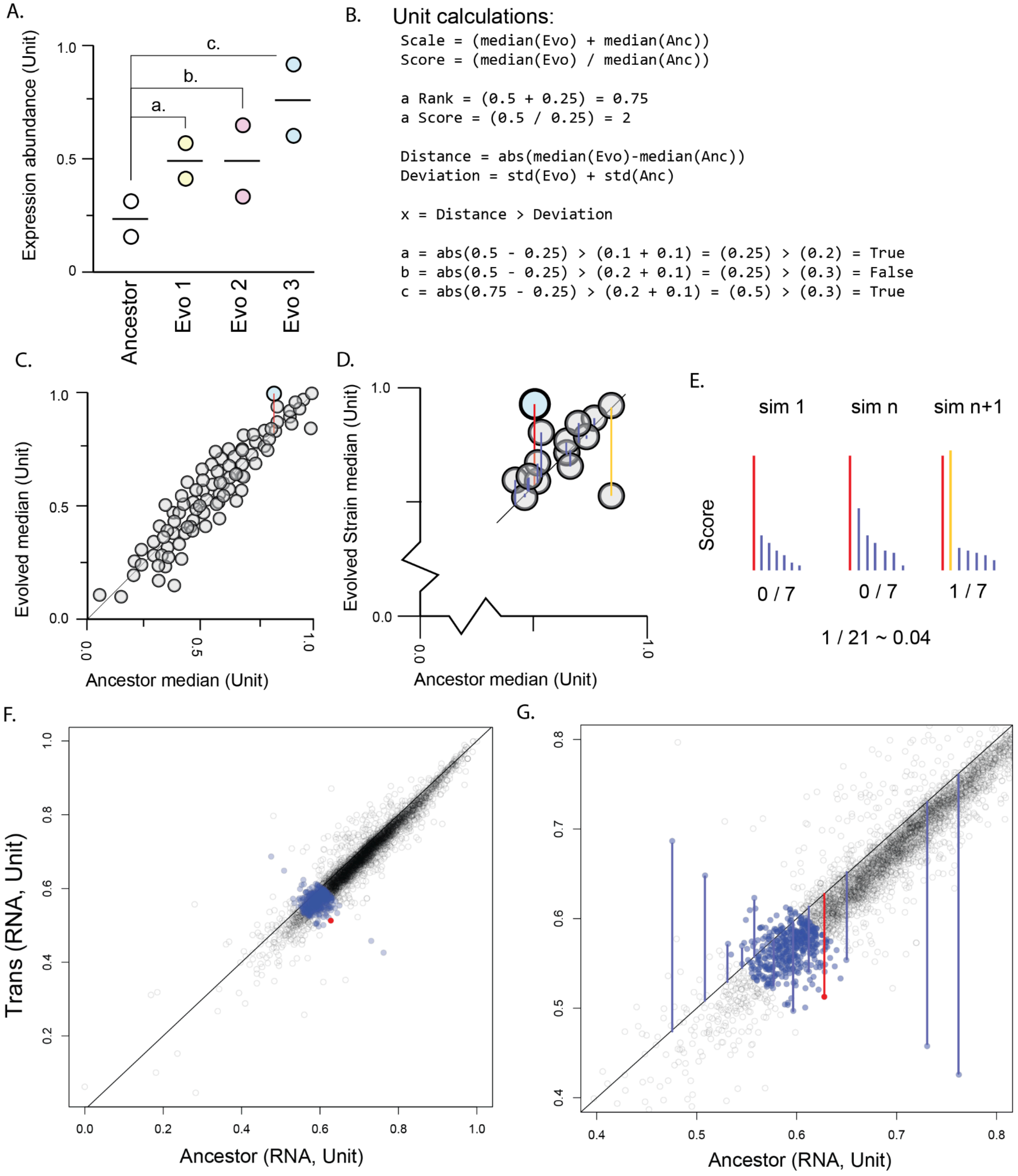


**Figure S6: Example of Unit DGEA at one level of expression**

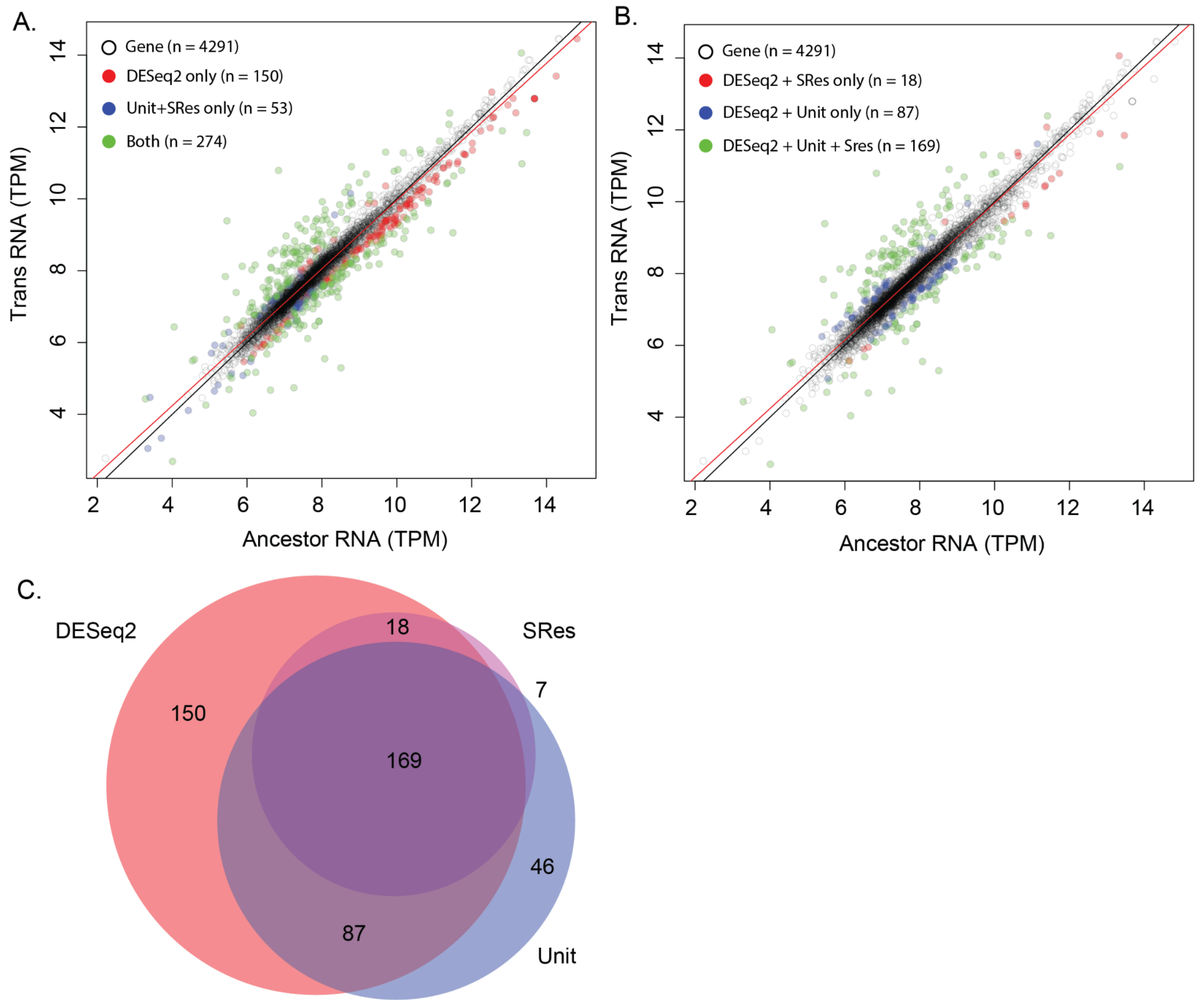


**Figure S7: Comparison of Unit+SRes analysis on differences in RNA abundance.** Performance compared to a DESeq2 test of RNA abundance finds reasonable agreement with a false positive rate of 0.01, a false negative rate of 0.35 and a F1 score of 0.73.

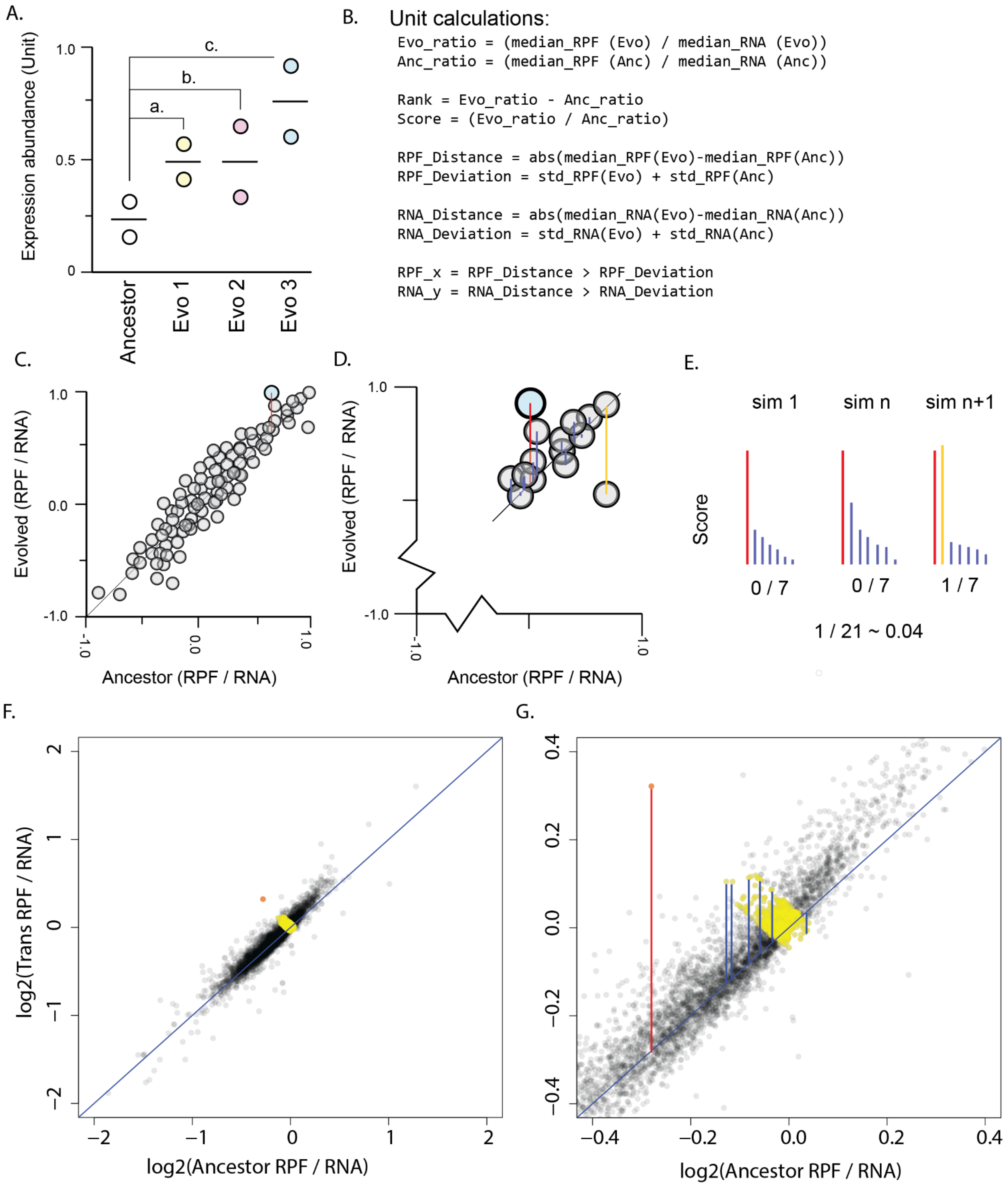


**Figure S8: Example of Unit analysis at two levels of expression**

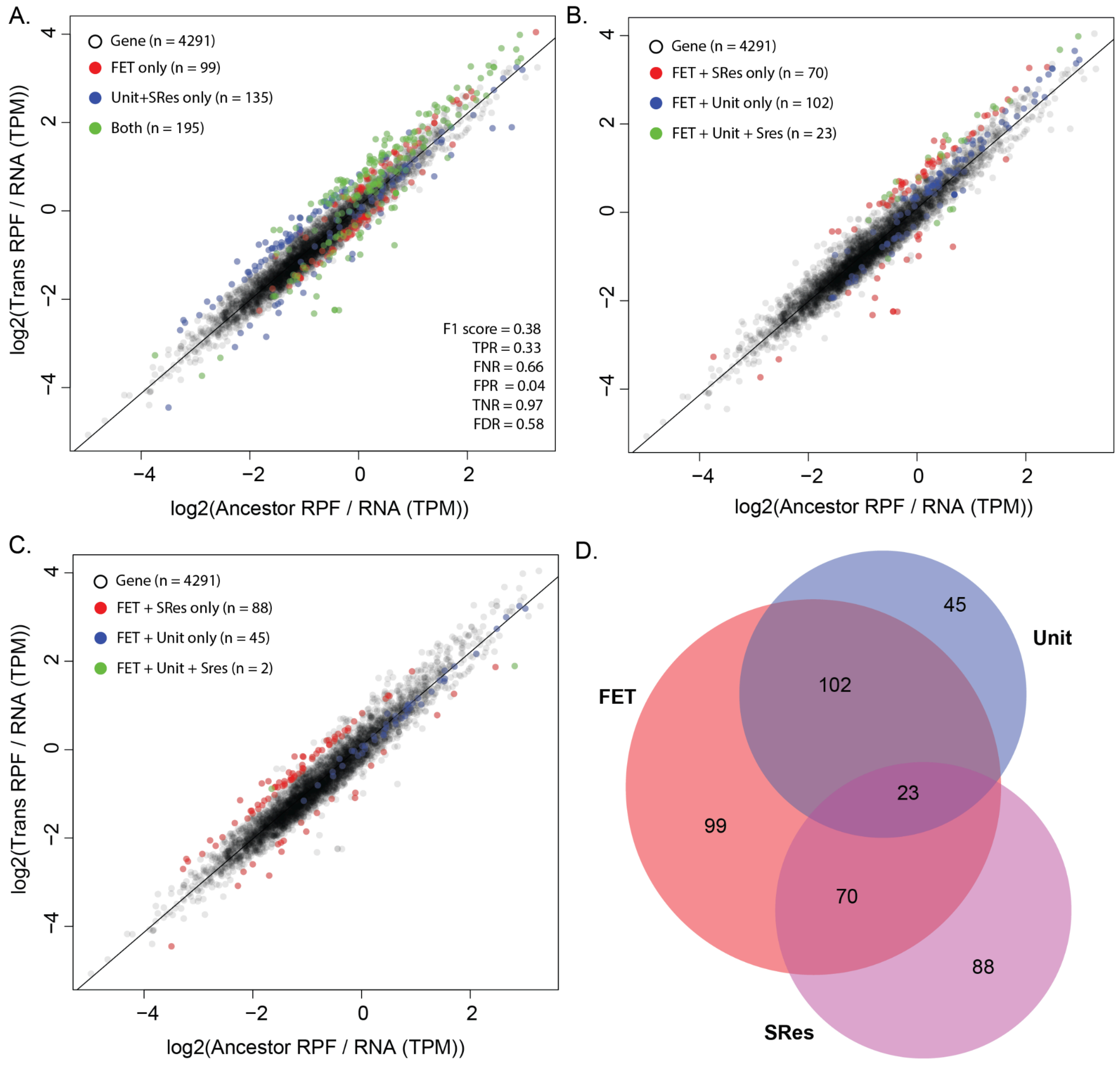


**Figure S9: Comparison of Unit+SRes analysis on differences in Translational Efficiency (RPF/ RNA).** Performance compared to FET test of translation efficiency ratios finds reasonable agreement with a false positive rate of 0.03, a false negative rate of 0.34 and a F1 score of 0.63.

### Supplemental tables:

**Supplemental table 1: ST1_chemostat_gene_relative_copy_number.tsv**

Table containing resolved copy number on a gene by gene by strain basis.

| **DGY1657** | RNA | RPF | MS |
|---|---|---|---|
| RNA | 0.98966 | 0.70425 | 0.584004 |
| RPF | | 0.990533 | 0.628283 |
| MS | | | 0.996504 |

| **DGY1726** | RNA | RPF | MS |
|---|---|---|---|
| RNA | 0.987811 | 0.635839 | 0.573157 |
| RPF | | 0.959884 | 0.60123 |
| MS | | | 0.998446 |

| **DGY1735** | RNA | RPF | MS |
|---|---|---|---|
| RNA | 0.987192 | 0.627072 | 0.545272 |
| RPF | | 0.979158 | 0.61208 |
| MS | | | 0.99717 |

| **DGY1741** | RNA | RPF | MS |
|---|---|---|---|
| RNA | 0.991494 | 0.612493 | 0.508551 |
| RPF | | 0.992748 | 0.582381 |
| MS | | | 0.989193 |

| **DGY1743** | RNA | RPF | MS |
|---|---|---|---|
| RNA | 0.982327 | 0.618318 | 0.50822 |
| RPF | | 0.987891 | 0.595538 |
| MS | | | 0.993212 |

**Supplementary Table ST2: Spearman *rho* correlation between replicates and between levels of expression.**

| **RNA** | DGY1726 | DGY1735 | DGY1741 | DGY1743 |
|---|---|---|---|---|
| DGY1657 | 0.963762 | 0.945636 | 0.885821 | 0.906666 |
| DGY1726 | | 0.978994 | 0.912389 | 0.939575 |
| DGY1735 | | | 0.904653 | 0.945145 |
| DGY1741 | | | | 0.939695 |

| **RPF** | DGY1726 | DGY1735 | DGY1741 | DGY1743 |
|---|---|---|---|---|
| DGY1657 | 0.946509 | 0.946274 | 0.892832 | 0.922275 |
| DGY1726 | | 0.9527 | 0.892116 | 0.939025 |
| DGY1735 | | | 0.876115 | 0.929458 |
| DGY1741 | | | | 0.916981 |

| **MS** | DGY1726 | DGY1735 | DGY1741 | DGY1743 |
|---|---|---|---|---|
| DGY1657 | 0.990968 | 0.990221 | 0.987052 | 0.989726 |
| DGY1726 | | 0.996515 | 0.991433 | 0.996051 |
| DGY1735 | | | 0.991952 | 0.996022 |
| DGY1741 | | | | 0.991495 |

**Supplementary Table ST3: Correlation between strains**

**Supplemental Table ST4: Results of DESeq2 on RNA**

Table containing the results of DESeq2 pairwise tests performed between each evolved strain and the ancestor using observed RNA abundances.

**Supplemental Table ST5: Results of DESeq2 on RPF**

Table containing the results of DESeq2 pairwise tests performed between each evolved strain and the ancestor using RPF abundances.

**Supplemental Table ST6: Results of Welch's t-test on MS intensities**

Table containing the results of Welch's t-test pairwise tests performed between each evolved strain and the ancestor using mass spectrometry instensities.

**Supplemental Table ST7: Genes with cluster assignment**

Table containing the gene name and assigned cluster for all genes in Supplemental Figure 3 and Figure 2A.

**Supplemental Table ST8: Results of DESeq2 on RNA Expected versus observed**

Table containing the results of DESeq2 pairwise tests performed between the expected RNA abundance of each evolved strain and the observed RNA abundance in the ancestor.

**Supplemental Table ST9: Results of Fisher's Exact Test on Translation Efficiency**

Table containing the results of Fisher's exact test conducted pairwise on ratios of TPM transformed RPF and TPM transformed RNA abundances for evolved strain and the ancestor.

**Supplemental Table ST10: Results of unit transform and test on MS and RPF**

Table containing the results of the unit transform and unit transform test. The test was conducted pairwise on ratios of unit transformed MS and unit transformed RPF abundances for evolved strain and the ancestor.

**Supplemental Table ST11: GATK Variant Calls**

Single nucleotide variants and indels identified by GATK in the strains in conjunction with variant effect prediction annotation by Ensembl VEP when possible. GATK was ran using default options against a modified reference genome containing the Gresham GFP reporter (Lauer et al. 2018). For Supplemental Figure 2 these were further reduced for the evolved by removing variants present in the ancestor.

**Supplemental Table ST12: CVish predicted boundaries**

CNV/SV boundaries for all strains predicted by CVish using Illumina sequencing reads. CVish was ran using default options against a modified reference genome containing the Gresham GFP reporter (Lauer et al. 2018). Each tab represents a strain and uses the gff format.

**Supplemental Table ST13: uORFish predicted uORF results.**

Table containing the results of uORFish ran on the ribosome profiling data for each replicate of each strain. uORFish was ran using default options, all potential predictions with a score of at least 0.5 are included. Only uORFs with scores of at least 0.9 in both replicates were included in the study.

**Supplemental Table ST14: Results of SSD1 motif scan**

Table containing the results of a motif search for the conserved *SSD1* motif ('CNYUCNYU', reported by Bayne et al. 2022) located within the transcript leaders (5'UTRs) of protein coding genes.

#### Supplemental files:

**Supplemental File 1: SF1_Saccharomyces_cerevisiae.R64-1-1.ncrna_wo_ncrna_genes.fa**

Custom fasta file used to filter out snRNA, snoRNA, rRNA, and tRNA but not potentially translated non-coding RNA.

**Supplemental File 2: SF2_MaxQuant_16plex_template_10262023.txt**

Isobaric weight template for MaxQuant for 16plex TMT-labeled MS.

**Supplemental File 3: SF3_parameters.txt**

Parameters file for MaxQuant analysis for MS data.
